# iGlu3Fast and iGlu3Slow, improved fluorescent reporters for detection of spontaneous glutamate activity in the brain

**DOI:** 10.1101/2024.08.30.610471

**Authors:** Oanh Tran, Sara Bertelli, Holly J. Hughes, Laura Rocha Llabrés, Tom Carter, Andrew J. R. Plested, Katalin Török

**Affiliations:** Neuroscience and Cell Biology Research Institute, City St George’s, University of London, London, UK; Humboldt Universität zu Berlin & Cluster of Excellence NeuroCure, Berlin, Germany

**Keywords:** biosensor, glutamate, neurotransmission, fluorescence, hippocampus

## Abstract

The genetically-encoded fluorescent glutamate sensor, iGluSnFR3, characterised by a high dynamic range and rapid *on*-kinetics, is an attractive sensor for glutamate imaging in the central nervous system. However, kinetic variants with ultrafast or slow *off*-kinetics are needed to broaden the spectrum of applications from monitoring rapid synaptic glutamate transients to mesoscale imaging of brain activity. Here we report binding-site variant S72T (iGlu3Fast) optimised for reporting fast glutamate release at individual sites with a fluorescence dynamic range of 57 and a decay *t*_1/2_ of 2 ms in solution, ∼5-fold faster than iGluSnFR3. In contrast, variants D25A and D25R (termed iGlu3Slow1 and iGlu3Slow2, respectively) presented with slow *off*-kinetics (decay *t*_1/2_ 43 ms and 30 ms, respectively, at 20°C), while retaining high dynamic ranges (48 and 65, respectively). These values were reduced when expressed in HEK293T cells, glutamate dissociation from iGlu3Fast slowed to a decay *t*_1/2_ of 6.9 ms, while decay *t*_1/2_-s of 115 and 234 ms were measured for iGlu3Slow1 and iGlu3Slow2, respectively. The slow decay rates in cells were in accordance with reduced *K*_d_-s in the range of 200 μM for iGlu3Fast and 6.4 and 8.4 μM for iGlu3Slow1 and iGlu3Slow2, respectively. 2-photon imaging in organotypic hippocampal slices reported spontaneous activity with high sensitivity by both iGlu3Fast and iGlu3Slow2. Strikingly, different kinds of glutamate transient were detected by the fast- and slow-decay sensors indicating that the rapid kinetics and increased fluorescence dynamic range make iGlu3Fast an excellent candidate for imaging high-frequency glutamate release at synapses while iGlu3Slow2 has the potential for mesoscale imaging of brain activity to record global events with high sensitivity.

## Introduction

Imaging neurotransmitter release at synapses represents a direct way of monitoring brain activity. The genetically-encoded glutamate sensor, iGluSnFR, opened a new era for imaging neuronal activity for glutamate, the most abundant excitatory neurotransmitter in the CNS (1, for a review see 2). While iGluSnFR represented a milestone in terms of visualising neurotransmitter release, its *off*-kinetics were fast enough to only resolve low frequency glutamate release. However, a binding site variant of iGluSnFR, iGlu*_u_* with ultrafast *off*-kinetics, enabled the resolution of high-frequency (100 Hz) glutamate release in hippocampal slices at individual CA3-CA1 synapses (3). iGlu*_u_* detected a reduced rate of glutamate clearance in the striatum in a mouse model of Huntington’s Disease (HD), revealing that in addition to diminished expression of EAAT2, transporter function is impaired in HD (4).

While further binding site mutations resulted in faster kinetics and/or reduced affinity of iGluSnFR (5), iGluSnFR3 has recently superseded iGluSnFR with increased dynamic range and faster on-kinetics (6). The design of iGluSnFR3 follows that of iGluSnFR, having the N and C-terminal portions of the bacterial GluBP fused to each terminus of a circularly permuted fluorescent protein, namely the more photostable cpSF-Venus. iGluSnFR3 has the same domain structure as iGluSnFR with PDGFR or other membrane targeting motifs used to target its expression to the extracellular face of the plasma membrane. iGluSnFR3 is the v857 variant of cpSF-Venus-based iGluSnFR that was developed by high-throughput technology incorporating several mutations, including the D25E mutation, previously used to modulate glutamate affinity and kinetics (3,5). The iGluSnFR3 v857 variant was selected for having the best fluorescence and kinetic properties of its generation (6). However, the kinetic limitations of iGluSnFR remain applicable to iGluSnFR3. Given the distinct advantages of a faster sensor like iGlu*_u_* for resolving glutamate release over a range of frequencies up to 100 Hz (3), we generated iGlu3Fast, an ultrafast variant of iGluSnFR3 with the S72T substitution, which adventitiously also further increased its fluorescence dynamic range. We furthermore generated two variants with slow *off*-kinetics (iGlu3Slow1 with the D25A and iGlu3Slow2 with the D25R mutation) expanding the scope towards mesoscale glutamate imaging.

## Materials and Methods

### Plasmids and cloning

Plasmid pCMV(MinDis).iGluSnFR (Addgene plasmid # 41732) and pRSET.GltI253-cpGFP.L1LV/L2NP (Addgene plasmid # 41733) were a gift from Loren Looger (1). Plasmids pRSET iGluSnFR3.v857 (Addgene plasmid # 175186), pAAV.hSyn.iGluSnFR3.v857.PDGFR (Addgene plasmid # 178329) and pAAV.hSyn.iGluSnFR3.v857.PDGFR-SGZ (Addgene plasmid #178330) were a gift from Kaspar Podgorski (6).

For expression in E. coli for protein purification, iGluSnFR3 and variants in the pRSET iGluSnFR3.v857 (Addgene plasmid # 175186) vector were used. For expression in HEK293T cells, the iGluSnFR3.v857 insert from Addgene plasmid # 178329 containing the IgK-chain, Myc epi tag and PDGFR TM domain was subcloned into the pEGFP-N1 vector replacing EGFP by restriction-ligation at the NheI and HindIII sites.

For expression in organotypic slices iGlu3Fast and iGlu3Slow2 were ligated into the pAAV shuttle vector (pAAV.hSyn.iGluSnFR3.v857.PDGFR-SGZ), replacing iGluSnFR3.v857 following digestion with BamHI and BstEII. AAV2/9 particles encoding iGluSnFR3, iGlu3Fast and iGlu3Slow2 were prepared by the Charité Viral Core Facility.

*E. coli* XL10-Gold and BL21 (DE3) Gold cells were purchased from Invitrogen.

### Site-directed mutagenesis

iGluSnFR3 variants in the pRSET vector were made using the Q5® Site-Directed Mutagenesis Kit (New England Biolabs) with the following primers (5’ - 3’):

iGluSnFR3 S72T (iGlu3Fast), TCC GAT TAC TAC GCA AAA CCG TAT TC iGluSnFR3 D25A (iGlu3Slow1), CGG TCA CCG TGC CTC TTC AGT GC iGluSnFR3 D25R (iGlu3Slow2), CGG TCA CCG TAG GTC TTC AGT GCC

iGluSnFR3 variants in the pEFP-N1 vector were generated using the following primers (5’ - 3’):

iGluSnFR3 S72T (iGlu3Fast), TCC TAT AAC AAC CCA AAA CCG CA iGluSnFR3 D25A (iGlu3Slow1), CGG CCA CCG AGC GTC TTC TGT GC iGluSnFR3 D25R (iGlu3Slow2), CGG CCA CCG AAG GTC TTC TGT GC

DNA sequencing was carried out by Genewiz.

### Protein expression and purification

iGlu3Fast, iGlu3Slow1, iGlu3Slow2 and iGluSnFR3.v857 (iGluSnFR3) in pRSET vector were overexpressed in *E. coli* BL21 Gold cells using 0.5 mM isopropyl thio-β-D-galactoside (IPTG) and purified on a NiNTA column (QIAGEN, ÄKTA Purifier, GE Healthcare) at 4 °C as described previously (3). Protein was checked for purity by SDS-PAGE then stored at –80°C.

### Protein concentration measurements

Protein concentration was calculated from the absorption spectra of the proteins at 280 nm using the molar extinction coefficient of 52830 M^-1^ cm^-1^ calculated using the ProtParam tool (Expasy).

### Equilibrium binding titrations

Fluorescence emission spectra were recorded using a Flurolog-3 spectrofluorimeter (Horiba) at excitation wavelength of 495 nm and emission wavelength of 518 nm, 20 °C. 250 nM iGlu3Fast, iGlu3Slow1, iGlu3Slow2 were titrated and compared with iGluSnFR3 in PBS buffer, pH 7.4 by stepwise addition of a solution of 50 - 500 mM glutamate until no further changes in fluorescence emission intensity were observed. Experiments were done in triplicates. Data was corrected for dilution, normalized and expressed as mean ± S.E.M. Apparent *K*_d_ values for L-glutamate, L-aspartate and L-glutamine (the ligand concentration at which half-maximum fluorescence intensity is achieved) were obtained by fitting the data to a single site binding equation using GraphPad Prism 9 software.

### Stopped-flow fluorimetry

Glutamate association and dissociation rates were measured using a TgK Scientific KinetAsyst™ double-mixing stopped-flow apparatus in single mixing mode. The fluorescence was excited at 495 nm and emission was collected using a 530 nm cut-off filter. For glutamate association, 250 nM iGlu3Fast, iGlu3Slow1, iGlu3Slow2 or iGluSnFR3 in PBS buffer, pH 7.4 was rapidly mixed with increasing concentrations of glutamate (from 100 – 500 mM stock solution of glutamate in PBS) at 20 °C. For glutamate dissociation, 250 nM protein in PBS buffer containing 100 - 300 µM glutamate was rapidly mixed with 1 - 3 mM purified bacterial periplasmic glutamate binding protein (GluBP) at 20 °C to quench glutamate released from the SnFR domain. Experiments were done at least in triplicates. Averaged data were fitted to a single exponential function using KinetAsyst software (TgK Scientific) to obtain association or dissociation rates. All concentrations given were the final concentrations in the mixing chamber.

### pH sensitivity

The apparent p*K*a was determined for iGlu3Fast and compared with iGluSnFR3, using a series of buffers. Depending on their respective pH buffering range, an appropriate buffer was used for the measurements (MES for pH 6-6.5, HEPES for pH 7-8, TRIS for pH 8.5-9 and CAPS for pH 10). The pH titrations were performed by recording fluorescence spectra in PBS only or PBS with saturating glutamate (30 mM) using 250 nM protein in 0.5 pH unit intervals at 20 °C (Fluorolog-3, Horiba). Excitation wavelength was set at 495 nm and emission spectra were run in the 500-600 nm range.

### Quantum yield determination

Measurements were made to compare iGlu3Fast and iGlu3Slow1 and iGlu3Slow2 with iGluSnFR3. Protein concentration was adjusted such that the absorbance at the excitation wavelength (492 nm) was between 0.001 and 0.1. A series of dilutions was prepared in PBS buffer only or PBS buffer with saturating glutamate (30 mM) and the fluorescence spectra were recorded on a Fluorolog-3 spectrofluorimeter (Horiba). For iGluSnFR3 and iGlu3Fast the quantum yield of GCaMP6f measured in Ca^2+^ -saturated buffer was used as a reference (Φ_+Ca2+_ = 0.59) (7). For iGlu3Slow1 and iGlu3Slow2 the quantum yield measured for Glu-bound iGlu3Fast of 0.94 was used. Data were plotted as integrated fluorescence intensity against absorbance and fitted to a linear regression (yielding the gradient, S). Quantum yields were obtained using the following equation: Φ_protein_ = Φ_reference_ × (S_protein_/S_reference_).

### Cell culture and transfection

For titration with glutamate HEK293T cells were grown in DMEM supplemented with 10% FBS and 4 mM L-Gln and maintained at 37°C in a 5% CO₂, at >100% humidity. Lipofectamine 2000 transfection reagent (Thermo Fisher Scientific) (2.5 μL per 35 mm glass bottomed dish) with pEGFP-N1 plasmids (2.5 μg per 35 mm glass bottomed dish) was used to introduce plasmids for the expression of iGluSnFR3 variants. Cells were imaged after 24 hours.

For patch-clamp fluorometry experiments HEK293T cells were grown in MEM supplemented with 6% FBS and maintained at 37°C in a 5% CO₂, 100% humidity atmosphere. Cells were transfected using PEI in a 1:3 ratio (vol/vol; DNA/PEI) the day after seeding. The mass ratio for co-transfection was 1:1 (GluA2 to each sensor pEGFP-N1-iGlu3Fast, pEGFP-N1-iGluSnFR3, pEGFP-N1-iGlu3Slow1, pEGFP-N1-iGlu3Slow2 or pCMV(MinDis).iGluSnFR), with a total of 2 μg DNA (1 μg/μL for each construct) per 35 mm dish. After 6 hours of incubation, the reaction was stopped, and the cells were split and seeded on 10 mm poly-lysine-coated coverslips. Patch-clamp fluorometry experiments were carried out 24–48 hours after transfection at room temperature.

### Wide field live cell fluorescence imaging

Live cell imaging was conducted using an Olympus IX71 inverted fluorescence microscope at room temperature. The imaging system comprised a monochromator (Optoscan, Cairn Research, Faversham, UK), a U-Plan-Apo x40 1.35NA objective and an Ixon3 EMCCD camera (Andor, Belfast, UK). Wavelength switching was synchronised with image capture using WinFluor software (Dr John Dempster, Strathclyde University). Glutamate sensors were illuminated with 488 ± 7 nm light. Excitation light was reflected onto the specimen using a 500DCXR dichrochic mirror (Chroma) and the emitted light passed through a 535 ± 50nm emission filter onto the chip of the EMCCD camera. The EMCCD camera was operated in frame transfer mode at full gain and cooled to −65°C. Full frame (512 by 512 pixels) images were acquired at 0.6 frames/sec in time lapse mode. Image sequences were analysed in ImageJ (https://imagej.net/ij/).

### Patch-clamp fluorometry of iGluSnFR3 variants expressed on HEK293T cells

The amplitude of the fluorescence response was determined by patch-clamp fluorometry as described previously (8). After obtaining whole-cell access, lifted cells were placed in front of a piezo-driven square barrel fast perfusion tool and subjected to the application of different solutions for activation by glutamate. Whole-cell patch clamp and simultaneous imaging recordings were performed in 158 mM NaCl, 20 mM HEPES-Na^+^ pH 7.4, 3 mM KCl and 1 mM CaCl₂ extracellular solution. The intracellular (pipette) solution contained 135 mM KCl, 20 mM KF, 20 mM HEPES-Na^+^ pH 7.4, 3 mM NaCl, 1 mM MgCl₂, and 2 mM EGTA at 23 °C (9,10). Currents from the co-expressed glutamate receptors were acquired using an Axopatch 200B amplifier and the Axograph acquisition program (Axograph Scientific) via an Instrutech ITC-18 D-A interface (HEKA Elektronik). After whole-cell configuration was obtained, cells were held at −50 mV, and current traces were filtered at 5 kHz and digitized at 20 kHz. Pipettes were made using borosilicate glass with an inner diameter of 1.17 mm (part no. G150TF-3 Multichannel Systems) and had a series resistance (Rₛ) between 2 and 6 MΩ, while the cell capacitance was between 5 and 25 pF, as measured by the compensating circuit of the amplifier. The maximum accepted voltage-clamp error, calculated as Iₘₐₓ * Rₛ, was 10 mV. The protocol consisted of repeated episodes in which cells underwent 300 ms piezo-controlled jumps to glutamate-containing solutions. The concentration-response analysis was performed by exposing cells to 1 μM, 10 μM, 100 μM, 500 μM, 1 mM, and 10 mM glutamate, applied in a sparse order. For imaging, an X-Cite Fluorescence Lamp was used for excitation light (Exciter 470/40 F49-470 AHF Analysentechnik) that was reflected onto the sample with a dichroic mirror (F48-495, AHF). Fluorescence emission passed through a 525/50 emission filter (F47-525) and was recorded at a frame rate of up to 200 Hz, without binning, on a Prime 95B CMOS camera (Photometrics). Camera acquisition was triggered directly from the digitizer. Image series were recorded with MicroManager (11) and analyzed with Fiji (12). We selected a region of interest (ROI) including the fluorescent cell and avoiding the patch pipette. All data were analyzed with IgorPro9 (Wavemetrics, Lake Oswego, OR, USA) and the Neuromatic plugin. To obtain the half maximal concentration of glutamate (*EC*_50_) and the maximum response (Δ*F*/*F*_0_ |_max_), we fit the concentration response data with a Langmuir binding isotherm:

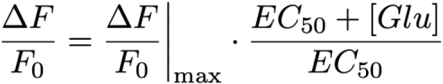

Decays were fit with a double exponential function:

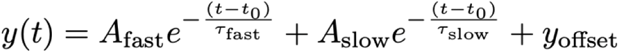

The offset was usually set to zero.

### Preparation of organotypic hippocampal slices

Organotypic hippocampal slices 350 μm thick, were obtained from 5- to 8-day-old (P5–P8) C57 black mice of either sex. Animals were sacrificed by decapitation, and the hippocampus was quickly resected and transferred into a cutting solution (13) containing (in mM): 0.5 CaCl₂, 2.5 KCl, 0.66 KH₂PO₄, 2 MgCl₂, 0.28 MgSO₄·7H₂O, 50 NaCl, 0.85 Na₂HPO₄·12H₂O, 25 glucose, 2.7 NaHCO₃, 175 sucrose, 2 HEPES, and 10 µg/mL red phenol, with a final osmolarity of 330 mOsm and with the pH adjusted to 7.3. The slices were prepared and plated on Millicell culture plates (EMD Millipore, Billerica, MA) according to the interface method (14,15), cultured in a MEM-based mouse slice culture medium, with the addition of 15% Horse Serum; 1x B27; 25 mM HEPES; 3 mM L-Glutamine; 2.8 mM CaCl_2_; 1.8 mM MgSO_4_; 0.25 mM Ascorbic Acid; 6.5 g/L D-Glucose and incubated at 34°C, 5% CO_2_ for up to 23 days, as previously described (16).

### Two photon imaging

Organotypic hippocampal slices were transduced with AAV2/9 encoding either iGluSnFR3.v857.SGZ (iGlusnFR3), AAV2/9.iGluSnFR3.v857.S72T.SGZ (iGlu3Fast), or AAV2/9.iGluSnFR3.v857.D25R.SGZ (iGlu3Slow2). For each slice, 5–10 µL of each AAV was applied at time points ranging from DIV 7 to DIV 14. No difference could be detected between slices transduced at early or late time points, so we pooled the data. The slices were imaged 6 to 10 days after AAV introduction (DIV 13–23). During recordings, slices were perfused with artificial cerebrospinal fluid (ACSF) solution containing (in mM) NaCl 145, KCl 2.5, HEPES 10, MgCl_2_ 1, CaCl_2_ 2, glucose 30. We found activity was almost non-existent at 23°C, the temperature often employed for organotypic slices, so the bath temperature was maintained at 34°C using a Temperature Controller TC02 (Multichannel Systems).

Fluorescence was excited with 940 nm illumination from a tunable Ti:Sapphire laser (Chameleon TPC, Coherent), scanned over the sample using a galvo-resonant scanner (8 kHz, custom Sutter BOB microscope built by Rapp Optoelectronic, Wedel, Germany). We imaged slices with either a 20x Zeiss objective (water immersion Plan-Apochromat with numerical aperture (NA) of 1.0 and working distance (WD) of 1.8 mm, 421452-9900) or a 63x Zeiss objective (water immersion Plan-Apochromat, NA = 1.0, WD = 2.1mm, 421480-9900). The tunable laser was routinely set to 30% of the full power (resulting in laser output of about 900 mW). Two-photon imaging was controlled using the ScanImage software package (Vidrio) and the laser beam was further attenuated (15-30% for 20x objective). For the 63x objective, which was substantially overfilled at the back aperture, we did not use further attenuation. The power at sample was typically 30 mW/mm^2^. Acquisition rates were 30 Hz for full frame and were increased to 70-200 Hz by selecting regions of interest. Typical pixel dwell times were 20-50 ns. For recording event kinetics, we selected regions of interest. Fluorescence emission was reflected by 565nm long-pass dichroic mirrors onto two photomultiplier tubes (H10770PA-40-04, Hamamatsu, Japan), one in the epi-configuration, and one coupled beneath the substage condenser. The photomultiplier outputs were combined, amplified by a DHCPA-100 transimpedance amplifier (at 80 MHz bandwidth; Femto, Germany) and digitised using a PXIe-1073 interface (National Instruments).

### Image data analysis

Image stacks obtained in WinFluor were imported into Image J for analysis. Background subtraction was performed to eliminate non-specific signals and images were converted from 8-bit to 32-bit format. Regions of interest (ROIs) corresponding to individual cells were defined by applying an intensity threshold. Pixels falling below the threshold were excluded from analysis by assigning them a ‘not a number’ (NaN) value. The average fluorescence intensity was then extracted. For each glutamate concentration the change in fluorescence Δ*F*/*F*_0_ was calculated.

Spontaneous activity from the 2-photon imaging time series was found either manually or using a workflow where the Z-projection of the standard deviation was divided by the Z-projection of the maximal intensity, followed by thresholding and mask generation. Time series were derived from a circular ROI around the identified spot. For calculation of fluorescence change (Δ*F*/*F*_0_), the baseline fluorescence (*F*_0_) was defined as the mean fluorescence in the period before the response and then subtracted from *j*th acquisition frame where the peak fluorescence was observed, and divided by *F*_0_ to convert each trace to units of Δ*F*/*F*_0_ according to equation: [Δ*F*/*F*]_j_=(*F*_j_−*F*_0_)/*F*_0_. For each ROI, we calculated a single Δ*F*/*F*_0_ value, corresponding to the maximal peak intensity observed during a transient for the particular movie. We measured *τ*_off_ by fitting the decay of each transient response. The rise time (10-90%) of individual transients was measured using the Neuromatic plugin (17).

## Results

### Biophysical characterisation of Fast and Slow iGluSnFR3 variants in solution

We generated the ultrafast variant iGluSnFR3 S72T (iGlu3Fast) and two slow variants D25A (iGlu3Slow1) and D25R (iGlu3Slow2) (**Figure 1**). Using purified protein, we compared their properties in solution with those of iGluSnFR3.

**Figure 1.**
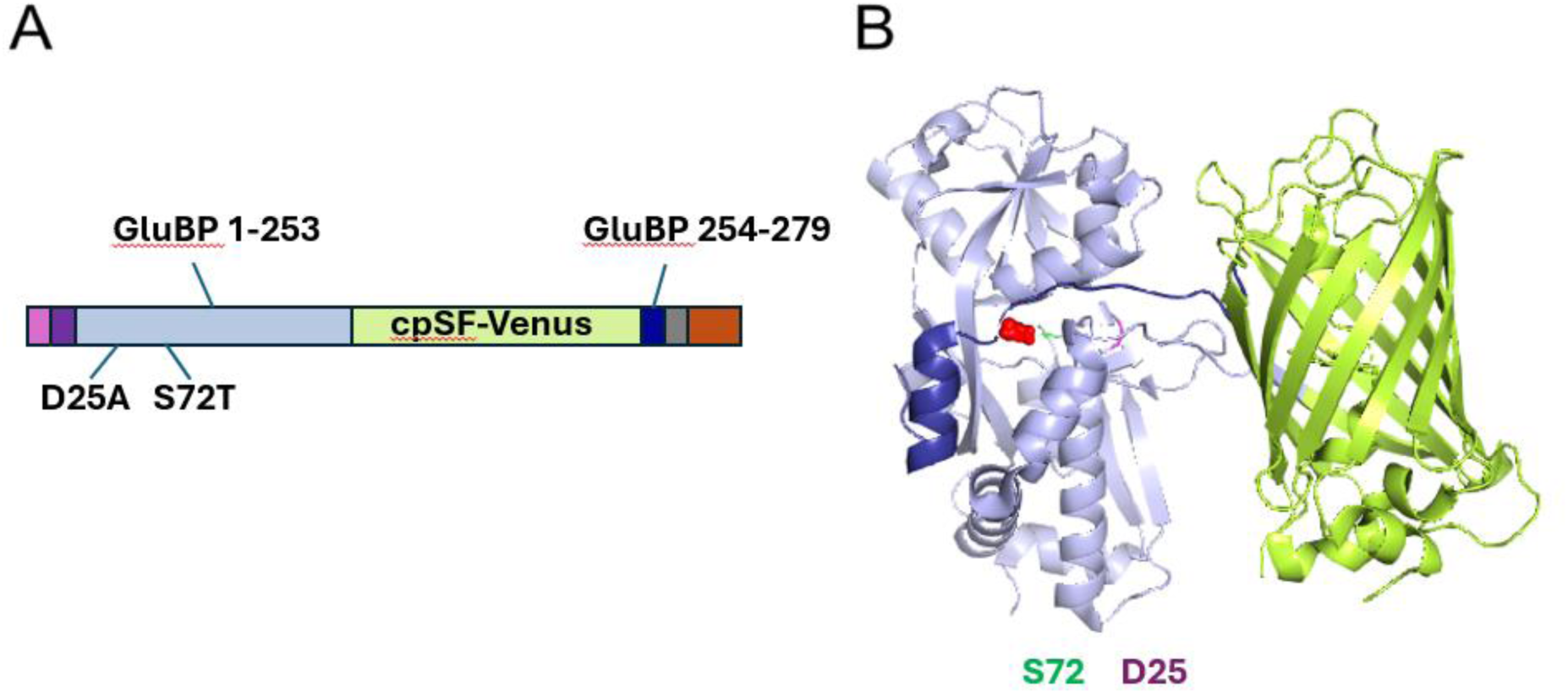
iGlu3Fast design. (**A**) Domain structure and design of single fluorophore-based genetically encoded glutamate indicator iGluSnFR3 indicating the S72T mutation; cpSF-Venus (yellow), IgG kappa secretion tag (pink), hemagglutinin (HA) tag (purple), myc tag (grey) and a PDGFR transmembrane domain (brown); GluBP 1-253 and 254-279 fragments are in light and dark blue, respectively. (**B**) Crystal structure of SF-iGluSnFR-S72A (8OVO) (18) with glutamate (red space filling display) incorporated to the binding site is used to illustrate the position of mutation sites D25 (red) and S72 (green).

First, we measured the fluorescence dynamic ranges of iGluSnFR3 and the variants. The iGlu3Fast variant superseded iGluSnFR3 with a relative fluorescence value of 57, compared to 48, measured for iGluSnFR3, in our hands (**Table 1** and **Supplementary Table 1**). The fluorescence dynamic ranges of iGlu3Slow1 and iGlu3Slow2 in solution were at 48 ± 1 and 65 ± 1, respectively (**Table 1** and **Supplementary Table 1**).

**Table 1.** Fluorescence dynamic range and kinetic parameters of iGluSnFR3 and variants in solution.

| Variant | <sup>a</sup>$F_{\max}/F_{\min}$ | $K_d$<br>mM | <sup>b</sup>$k_{-2}$<br>s<sup>-1</sup> | $K_1$<br>mM<sup>-1</sup> | $k_{+2}$<br>s<sup>-1</sup> | <sup>c</sup>$K_d$<br>(calc)<br>mM |
| --- | --- | --- | --- | --- | --- | --- |
| iGlu3Fast | 57 ± 1 | 3.18 ± 0.051 | 340 ± 33 | 0.074 ± 0.035 | 832 ± 147 | 3.62 |
| iGluSnFR3 | 48 ± 1 | 0.373 ± 0.003 | 71 ± 3 | 0.111 ± 0.011 | 1399 ± 43 | 0.398 |
| iGlu3Slow1 | 48 ± 1 | 0.058 ± 0.001 | <sup>a</sup>16 | 0.111 ± 0.007 | 1260 ± 34 | 0.115 |
| iGlu3Slow2 | 65 ± 1 | 0.204 ± 0.008 | <sup>a</sup>23 | 0.041 ± 0.010 | 1736 ± 278 | 0.319 |

$k_{+2}$ , $k_{-2}$ and $K_1$ are fitted values to Scheme 1 as described in (3). <sup>a</sup> $F_{\max}/F_{\min}$ is the ratio of fluorescence intensities in the presence and absence of saturating glutamate. <sup>b</sup>The measured *off*-rates corresponded to the dissociation rate constants and were thus used to constrain the hyperbolic fit. $K_d$ was measured by equilibrium binding. <sup>c</sup> $K_d$ (calc) is the value calculated from the fitted kinetic constants.

Second, we determined the *off*-kinetics of iGluSnFR3 and the variants. Using excess GluBP to trap the released glutamate, the rate constants for glutamate dissociation were 340 ± 48 s^-1^ (*t*_1/2_ 2 ms) and 71 ± 3 s^-1^ (*t*_1/2_ 10 ms) for iGlu3Fast and iGluSnFR3, respectively, at 20 °C (**Figure 2A&B**). For iGlu3Slow1 and iGlu3Slow2, the *off*-rate constants were 16 s^-1^ (t_1/2_ 43 ms) and 23 s^-1^ (t_1/2_ 30 ms), respectively (**Figure 3A&B**).

**Figure 2.**
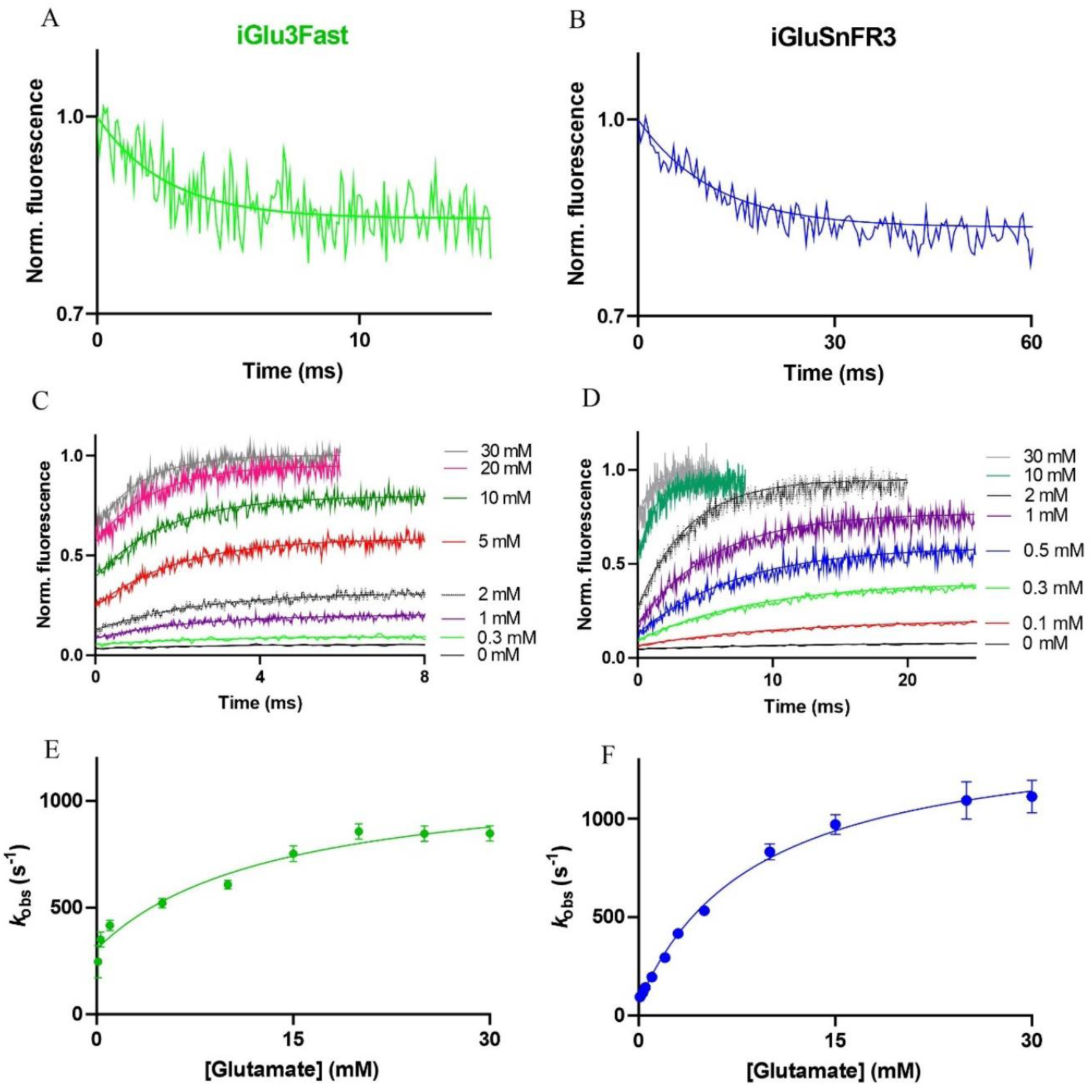
Kinetics of glutamate binding by iGluSnFR3 and its fast decay variant iGlu3Fast Glutamate dissociation kinetics (**A&B)** and glutamate association kinetics (**C&D**) of iGlu3Fast and iGluSnFR3. Stopped-flow records of iGlu3Fast and iGluSnFR3 fluorescence during the reaction with the indicated concentrations of glutamate. Experimental data are overlaid with curves fitted to single exponentials; (**E&F**) Plot of observed association rates, *k*_obs(on)_ of iGlu3Fast and iGluSnFR3 as a function of glutamate concentration. All experiments were carried out at 20°C.

**Figure 3.**
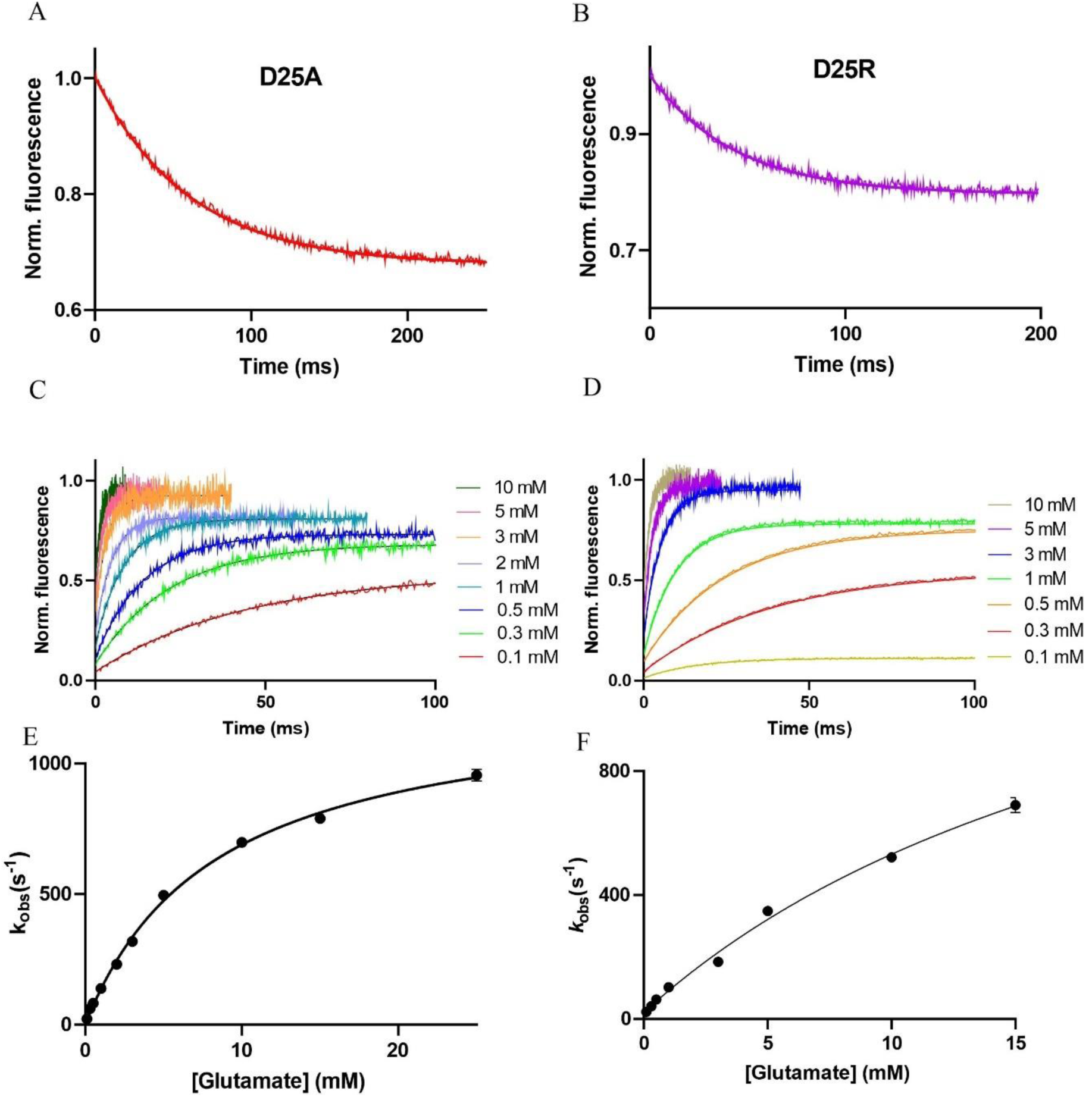
Kinetics of glutamate binding by iGluSnFR3 slow decay variants iGlu3Slow1 and iGlu3Slow2. Glutamate dissociation kinetics of (**A**) iGlu3Slow1 (D25A) and (**B**) iGlu3Slow2 (D25R) determined by stopped-flow fluorimetry at 20°C. Experimental data are overlaid with curves fitted to single exponentials. Fluorescence changes are normalised to *F*_max_ of 1. Glutamate association kinetics of (**C**) iGlu3Slow1 and (**D**) iGlu3Slow2. Stopped-flow records show reactions with the indicated concentrations of glutamate. Experimental data are overlaid with curves fitted to single exponentials; (**E&F**) Plot of observed association rates, *k*_obs(on)_ of iGlu3Slow1 and iGlu3Slow2 as a function of glutamate concentration.

We next determined the glutamate concentration dependence of the *on*-kinetics by rapidly mixing the sensor protein with different concentrations of glutamate (**Figure 2C&D, 3C&D)**.

The plots of the observed rates against the glutamate concentration had a hyperbolic shape (**Figure 2E&F, 3E&F)**. The data were consistent with the mechanism shown in **Figure 4**. The rate plots were fitted to a hyperbolic function that describes a two-step reaction, in which rapid binding is followed by a conformational change and in which fluorescence enhancement occurs in the second step, as found previously for iGluSnFR and its ultrafast variant iGlu*_u_* (3). Fitting the data to the equation that provides analytical solution for the mechanism (3). From that the equilibrium binding constant for the first step as well as the forward and reverse rate constants for the second, isomerization step (*K*_1_, *k*_+2_, *k*_-2_, respectively) were determined. **Table 1** shows all the fitted parameter values for the reaction scheme. Best fit values for the rate limiting *on*-rate (*k*_+2_) were 832 ± 148 s^-1^ and 1399 ± 43 s^-1^, 1260 ± 34 s^-1^ and 1736 ± 278 s^-1^ for iGlu3Fast, iGluSnFR3, iGlu3Slow1 and iGlu3Slow2, respectively, showing that the variants retained the rapid *on*-response of iGluSnFR3.

**Figure 4.**
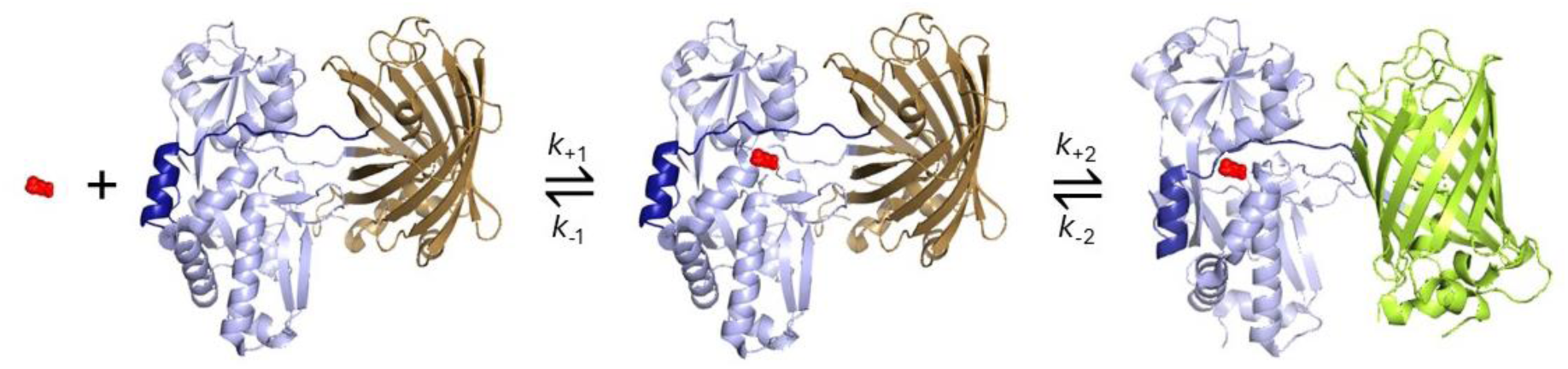
Two-step model of the kinetic mechanism of genetically encoded glutamate sensors. Cartoon diagram depicting the putative molecular transitions of iGlu3Fast and iGluSnFR3 variants to the fluorescent state. Key: GluBP 1-253 (light blue) and 254-279 (dark blue) fragments, cpSF-Venus (sand and limon), glutamate (red). The apo-state and initial Glu-bound state are modelled on the structure of the SF-iGluSnFR-S72A in complex with citrate (PDB ID 8OVN (18) as an approximation as citrate binding to GluBP does not evoke a fluorescence enhancement (19). The high-fluorescence state is modelled on the structure of the SF-iGluSnFR-S72A in complex with aspartate (PDB ID 8OVO (18). cpSF-Venus is dim (sand) and remains dim upon Glu binding until a conformational change involving both GluBP and cpSF-Venus leads to an enhancement of fluorescence emission. In GluPB the closure of the clamshell stabilizes its structure forcing a reorientation of cpSF-Venus relative to GluBP whereby the linker regions seal the cpSF-Venus β-barrel (limon).

Dissociation constants (*K*_d_) for glutamate binding to purified proteins were measured by equilibrium titration in solution. Strikingly, the *K*_d_ for iGlu3Fast was 3.18 ± 0.05 mM, 8-fold higher than iGluSnFR3 (373 ± 3 μM) (**Figure 5A**, **Table 1**). *K*_d_ values for iGlu3Slow1 and iGlu3Slow2 were 58 ± 1 μM and 204 ± 8 μM, respectively (**Figure 5A**, **Table 1**), 6.4- and 1.8-reduced compared to iGluSnFR3. The *K*_d_ values calculated from the fitted constants were in good agreement with those measured by equilibrium titration (**Table 1**).

**Figure 5.**
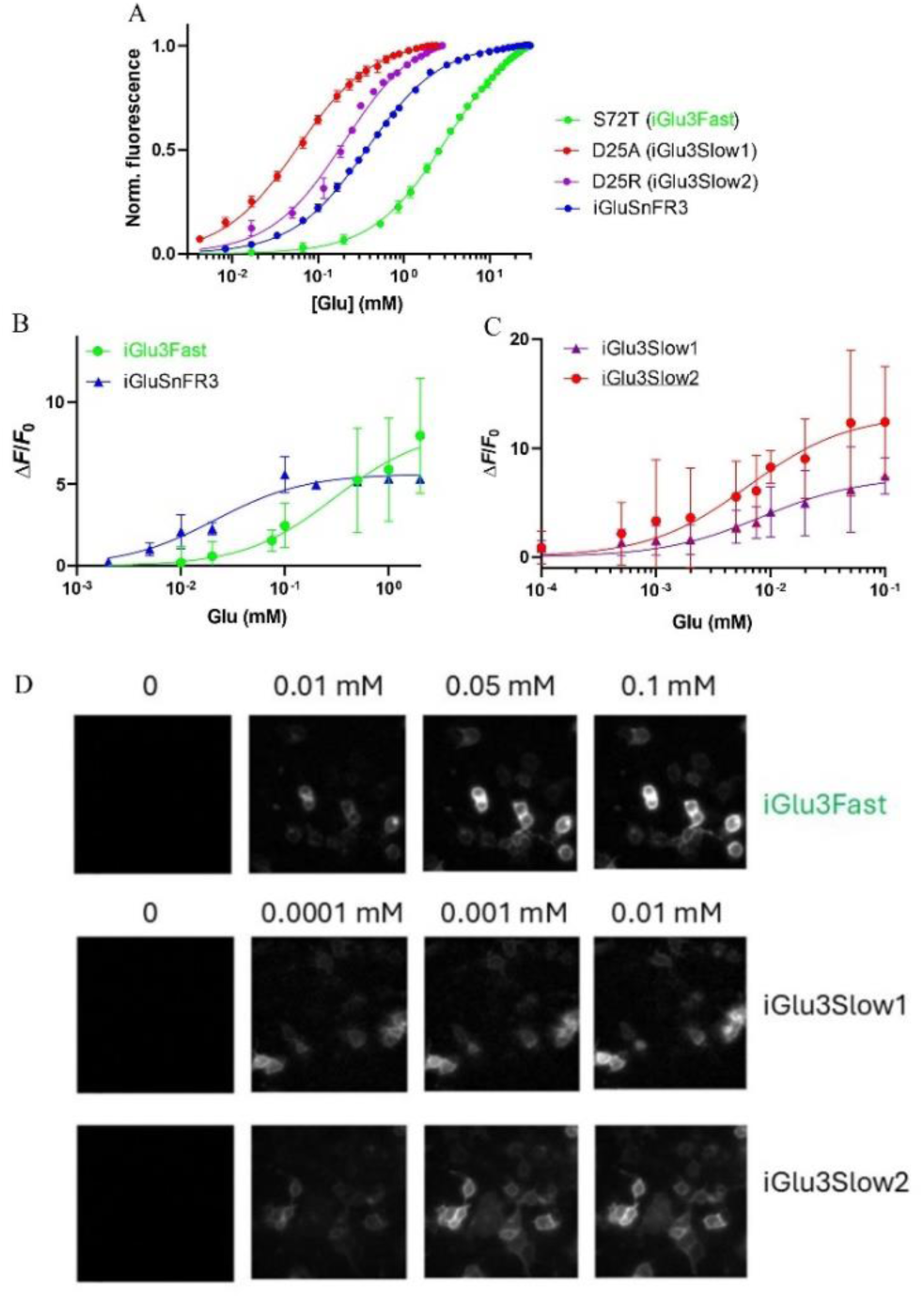
Apparent affinities of iGluSnFR3 and variants for glutamate in solution and in HEK293T cells. (**A**) Equilibrium binding titration of iGluSnFR3 and variants in solution. (**B**) Glutamate equilibrium binding titration of iGlu3Fast and iGluSnFR3 expressed on the surface of HEK293T cells. (**C**) Glutamate equilibrium binding titration of iGlu3Slow1 and iGlu3Slow2 expressed facing the extracellular milieu. (**D**) Representative images of HEK293T cells expressing the iGluSnFR3 variants titrated with glutamate. Glutamate concentrations in selected samples given.

### Brightness and quantum yield measurement of iGlu3Fast

Brightness and quantum yield data for iGluSnFR showed that the increase in brightness upon glutamate binding was almost entirely accounted for by an ∼4-fold increase in the extinction coefficient (3). In contrast, for iGluSnFR3, increases seen in both the extinction coefficient and the quantum yield upon glutamate binding agree with previous reporting (6). Similarly, the increased brightness of glutamate-bound iGlu3Fast and iGlu3Slow1 derived from increases in both the extinction coefficient and the quantum yield with overall increased brightness and fluorescence dynamic range (**Table 1** and **Supplementary Table 1**). Because of the low fluorescence of the apo-forms, it is challenging to measure both the extinction coefficient and the quantum yield accurately. In summary, the greater dynamic range of iGluSnFR3 and the further increased dynamic range of iGlu3Fast derive from the lower brightness of their apo-forms together with the large increases in their quantum yield upon glutamate binding, rather than greater brightness of their glutamate-bound forms (**Supplementary Table 1**).

### Ligand selectivity and pH dependencies of iGlu3Fast and iGlu3Slow2

The selectivity patterns of iGlu3Fast and iGlu3Slow2 with respect to glutamine were similar to that of iGluSnFR3 with the fluorescence enhancement following glutamine binding being barely detectable. Fluorescence enhancement by aspartate binding for iGlu3Fast and iGlu3Slow2 was ∼4- and 2-fold reduced, respectively, compared to that by glutamate (**Supplementary Figure 1**, **Supplementary Table 2**). Consistent with a previous report for the S72T variant of iGluSnFR3 (6), iGlu3Fast showed a notable increase in aspartate affinity. Measurement of the pH dependence of the fluorescence enhancement of the Glu-bound form of iGlu3Fast and iGlu3Slow2 revealed a p*K*_A_ value of 7 for both compared to 7.5 for iGluSnFR3. The pH ranges in which the Δ*F*/*F*_o_ signal remained stable were 7-8 for iGlu3Fast, 6.5-7.5 for iGlu3Slow2 and 6-7.5 for iGluSnFR3 (**Supplementary Figure 2**).

### Glutamate affinity of iGluSnFR3 variants expressed in HEK293T cells

The glutamate affinity of iGluSnFR and its variants e.g. iGlu*_u_* were found to increase when the sensor protein is expressed in cells facing the extracellular space (1,3,4). Although the mechanism is unknown, we also expected that the fluorescence dynamic ranges would be reduced by the cellular environment. We therefore determined the apparent affinities (*K*_d_) and dynamic ranges of the iGluSnFR3 variants in HEK293T cells in live cell imaging experiments by equilibrium titration with glutamate over a range of concentrations (**Figure 5B-D**). The measured values are shown in **Table 2**. The Δ*F*/*F*_0_ values were reduced up to 5-fold following expression in mammalian cells. The *K*_d_ values were more strongly affected exhibiting up to 25-fold reduction, indicating tighter binding for the cell-surface sensors.

**Table 2.** Apparent *K*_d_ values and dynamic ranges of the iGluSnFR3 variants determined by equilibrium titration with glutamate in HEK293T cells.

| Variant | $K_d$ (μM) | $R1$ | $\Delta F/F_0$ | $R2$ |
| --- | --- | --- | --- | --- |
| iGlu3Fast | $302 \pm 107$ | 10.5 | $8.3 \pm 0.9$ | 5.1 |
| iGluSnFR3 | $21 \pm 4$ | 3.4 | $5.6 \pm 0.3$ | 4.6 |
| iGlu3Slow1 | $6.4 \pm 1.4$ | 9.1 | $13.1 \pm 0.9$ | 2.5 |
| iGlu3Slow2 | $8.4 \pm 2.1$ | 24 | $7.4 \pm 0.6$ | 3.1 |

The ratios *R*_1_ and *R*_2_ represent the factors by which apparent *K*_d_ and Δ*F*/*F*_0_ values, respectively, measured in solution were decreased when the variants were expressed on the cell surface.

### In situ kinetics of iGluSnFR3 variants determined by patch-clamp fluorimetry

We next compared iGlu3Fast, iGlu3Slow1 and iGlu3Slow2 kinetics in situ expressed in HEK293T cells using patch-clamp fluorometry (20). Cells were co-transfected with AMPA receptor subunit GluA2 to verify glutamate solution application by measuring glutamate-mediated currents from the AMPA receptors whilst simultaneously recording fluorescence dynamics of the glutamate sensors in response to glutamate application. This approach served as a control for both the speed of rapid perfusion switching and cell health during concentration-response experiments.

Both iGlu3Fast and iGluSnFR3 responded stably to jumps into 1 mM glutamate with fast activation (**Figure 6A**). The decay after glutamate removal was faster for iGlu3Fast (see below). When expressed on the membrane of HEK293T cells, iGlu3Slow1 and iGlu3Slow2 exhibited much slower decay kinetics than iGlu3Fast and iGluSnFR3, requiring a modified acquisition protocol with shorter jumps into glutamate solution and extended intervals between applications to permit accurate measurements of the decay. However, the robust response to 20 ms jumps into 1 mM glutamate remained highly reproducible (**Figure 6B&C**).

**Figure 6.**
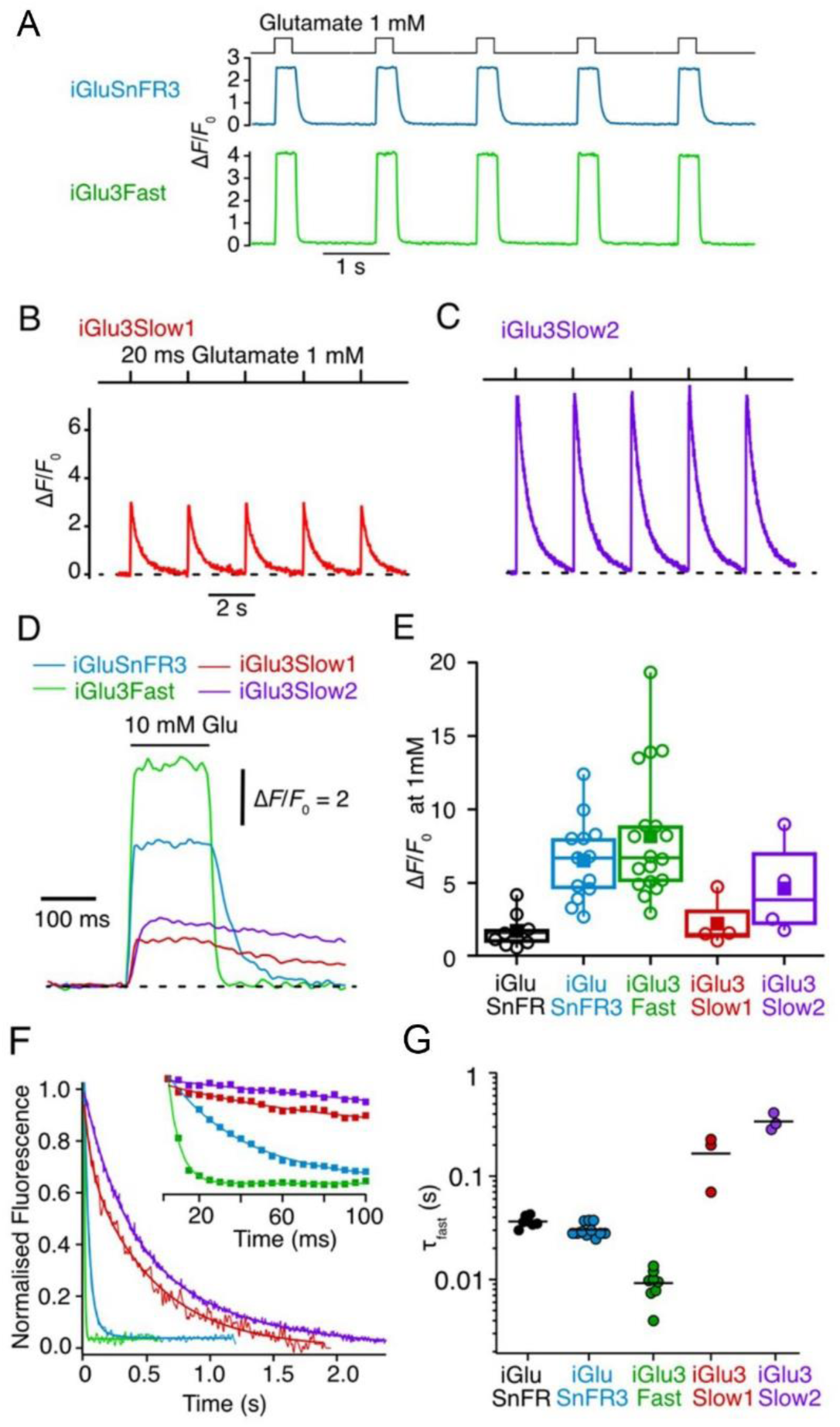
The fractional fluorescence changes and decay kinetics of iGluSnFR variants in HEKT293 cells. (**A**) Trains of consecutive fluorescence responses for membrane-expressed iGluSnFR3 and iGlu3Fast in whole lifted cells to 200 ms jumps into 1 mM glutamate. Responses of (**B**) iGlu3Slow1 and (**C**) iGlu3Slow2 measured by patch-clamp fluorometry. Glutamate concentration was jumped to 1 mM for 20 ms. Images were recorded at 200 frames/s. (**D**) Representative traces displaying the fluorescence increase (Δ*F*/*F*_0_) and response kinetics of all iGluSnFR3 variants to 10 mM glutamate. (**E**) Average fractional fluorescence changes for all the variants measured upon 1 mM glutamate application. Mean values were as follows: 1.7 ± 1.1 (iGluSnFR, *n* = 9), 6.5 ± 2.7 (iGluSnFR3, *n* = 13), 8.2 ± 4.1 (iGlu3Fast, *n* = 18), 2.3 ± 1.4 (iGlu3Slow1, *n* = 4), and 4.6 ± 2.8 (iGlu3Slow2, *n* = 4), respectively. (**F**) Representative traces of the decay kinetics after a brief jump into 1 mM glutamate solution. (**G**) Summary of the fast decay time constants (*τ*_fast_) for all iGluSnFR3 variants and iGluSnFR. Mean *τ*_fast_ values: 36 ± 4 ms (iGluSnFR), 30 ± 4.3 ms (iGluSnFR3), 9.2 ± 2.7 ms (iGlu3Fast), 166 ± 69 ms (iGlu3Slow1), and 339 ± 53 ms (iGlu3Slow2). Although best fits to the date were obtained by using a double exponential function, normally ∼98% of the amplitude represented the fast component, hence *τ*_fast_ values are reported.

At 1 mM glutamate, the response of iGlu3Fast expressed on the cell surface displayed a greater dynamic range than iGluSnFR3 (8.2 vs. 6.5; **Figure 6C&D**). At this concentration, iGluSnFR3 is already saturated. Upon exposure to 10 mM glutamate, the greater dynamic range of iGlu3Fast became even more evident (**Figure 6D**). The dynamic range of iGlu3Slow2 in response to 1 mM glutamate was substantially greater than that of iGlu3Slow1 (4.6 vs 2.3; **Figure 6C&D**).

The limit of solution exchange at a whole lifted cell is in the range 10-20 ms (21). When measuring the *off*-rate for the fluorescent signal for iGlu3Fast, best fits to the data were obtained by using a double exponential function. However, as ∼98% of the amplitude represented the fast component, *τ*_fast_ values are reported to describe the signal decay. *τ*_fast_ for iGlu3Fast was ∼4-fold faster than for iGluSnFR3, with a mean *τ*_fast_ of 9.2 ms (**Figure 6F&G**). This is similar to the decay of the AMPA receptor current in the same conditions, and at the limit of our recording system and technique (200 Hz image collection) making the response to 10 mM glutamate almost square. On the other hand, the decay time constants for iGlu3Slow1 and iGlu3Slow2 were 166 ms and 340 ms, corresponding to 5- and 9-fold slower decay times than iGluSnFR3, respectively **(Figure 6G)**.

Interestingly, the slow variants also showed slower apparent *on*-kinetics (**Figure 6C**), although a quantitative comparison with faster variants was limited by sampling rates for the patch clamp fluorometry system (up to 200 Hz).

For any receptor with fast kinetics, it is well known that a fast concentration jump (non-equilibrium measurement) can give a different concentration-response relation from an equilibrium titration (22). Therefore, the apparent glutamate affinities of iGluSnFR3 and iGlu3Fast were compared with fast solution exchange concentration-response analysis. We performed a dose-response analysis to compare glutamate affinities of iGluSnFR3 and iGlu3Fast. Representative images of cell fluorescence across exposure to increasing glutamate concentrations (**Figure 7A**) illustrate a higher *K*_d_ for iGlu3Fast, with an *EC*_50_ of 148 ± 68 µM, approximately 10-fold higher than iGluSnFR3 (12 ± 5 µM) (**Figure 7B**), in reasonable agreement with the cell titration assay presented in **Figures 5B&C**, and consistent with previous findings for iGluSnFR variants (1,3,6). Notably, although the extent of reduced affinity of iGlu3Fast for glutamate over iGluSnFR3 was maintained, while both iGlu3Fast and iGluSnFR3 showed ∼25-fold higher affinity for glutamate in cells compared to purified proteins in solution **(Figure 5A)**.

**Figure 7.**
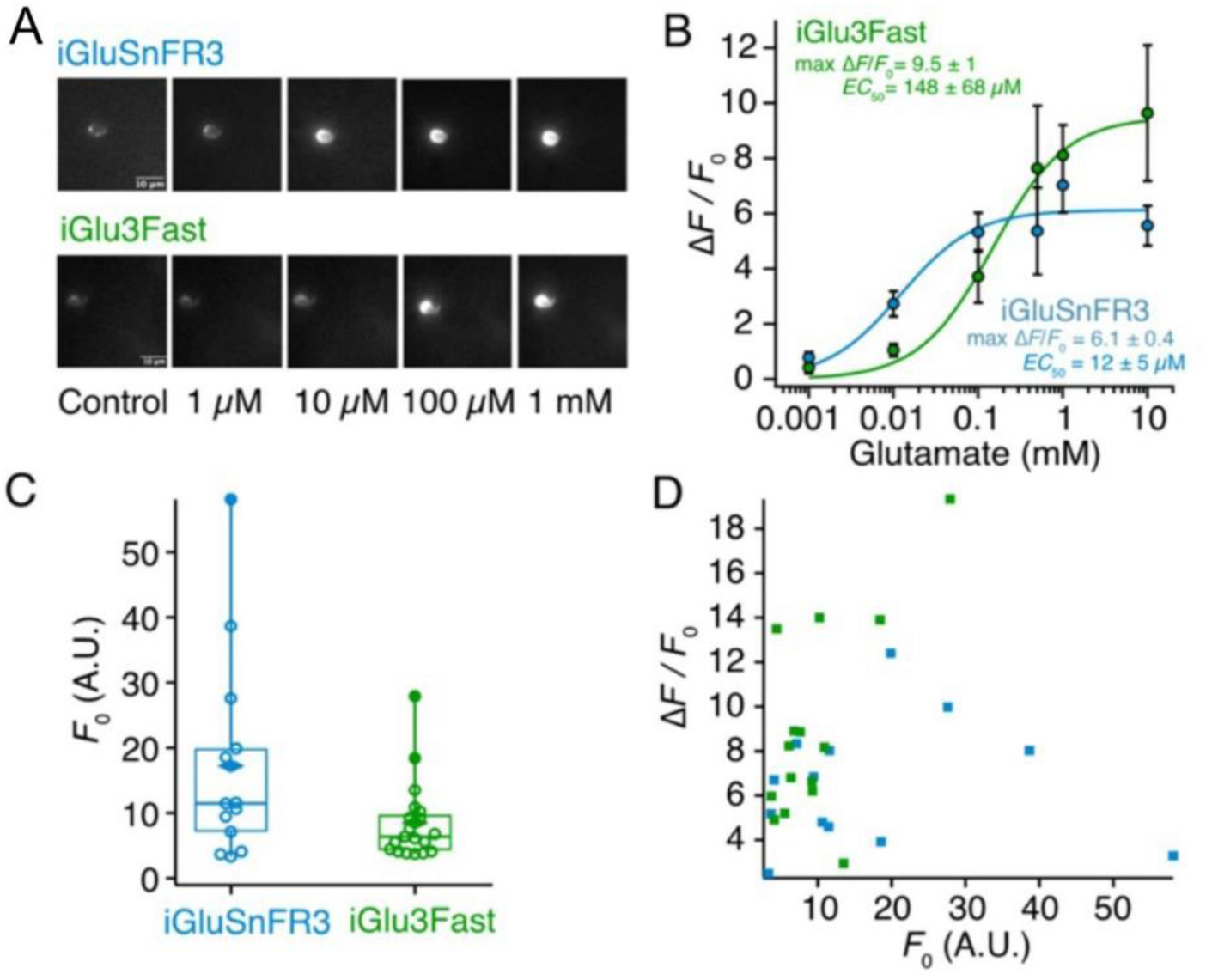
Concentration-response analysis of iGluSnFR3 and iGlu3Fast in HEK293 cells. **(A)** Representative images of the baseline fluorescence (*F*_0_, Control) and glutamate-induced fluorescence during 200 ms jumps into increasing glutamate concentrations (1 µM, 10 µM, 100 µM, 1 mM). Each montage is for a single lifted cell expressing iGluSnFR3 (upper), or iGlu3Fast (lower). (**B**) Concentration-response curve for iGluSnFR3 (blue) and iGlu3Fast (green). Fitted maximum responses (Δ*F*/*F*_0_) were 6.1 ± 0.4 (iGluSnFR3) and 9.5 ± 1 (iGlu3Fast). The half maximal concentrations (*EC*_50_) from the fits were 12 ± 5 µM (iGluSnFR) and 148 µM ± 68 µM (iGlu3Fast). (**C**) Box plot of the baseline cell fluorescence (*F*_0_) before application of glutamate. Diamonds represent mean values. (**D**) Scatter plot of relation between baseline fluorescence (*F*_0_) and the corresponding fold change increase upon 1 mM glutamate application (Δ*F*/*F*_0_, dynamic range). Dots represent individual cells.

The basal fluorescence in iGlu3Fast-transfected cells appeared lower than with iGluSnFR3, but this effect was weak because of the wide range of values (**Figure 7D**). Nevertheless, we evaluated whether differences in basal fluorescence could explain the enhanced dynamic range of iGlu3Fast. Overall, there was no correlation between the basal fluorescence and the observed dynamic range for either iGluSnFR3 or iGlu3Fast (**Figure 7E**).

In summary, iGlu3Fast stands out with a greater dynamic range, ∼5-fold faster *off*-rate and a 10-fold higher *K*_d_ for glutamate when displayed on the plasma membrane of HEK293T cells, representing the sensor most suitable in the field for monitoring synaptic release with high temporal resolution. The iGlu3Slow1 and iGlu3Slow2 variants offer high sensitivity and long-lasting fluorescence signals for large-scale imaging.

### iGluSnFR3 variants report spontaneous synaptic glutamate release in hippocampal organotypic slice by two-photon imaging

The primary goal of generating new variants of iGluSnFR is to improve the reporting of glutamatergic neurotransmission. To provide a reference for recording glutamate transients in a native environment, we expressed iGluSnFR3 and variants in mouse organotypic hippocampal slices by transducing them with AAVs, and recorded fluorescence signals from two photon excitation at up to 200 Hz frame rate (for cropped regions of interest).

The iGluSnFR3 sensor displayed a diffuse baseline signal in two-photon imaging when expressed in organotypic hippocampal slices, with spontaneous glutamate release detectable across multiple regions, including the dentate gyrus, CA1, and CA3. Similar to previous reports, the decay τ of spontaneous events was >100 ms (**Figure 8A**). We observed that the spatial extent of the fluorescence was relatively broad (see **Figure 9**).

**Figure 8.**
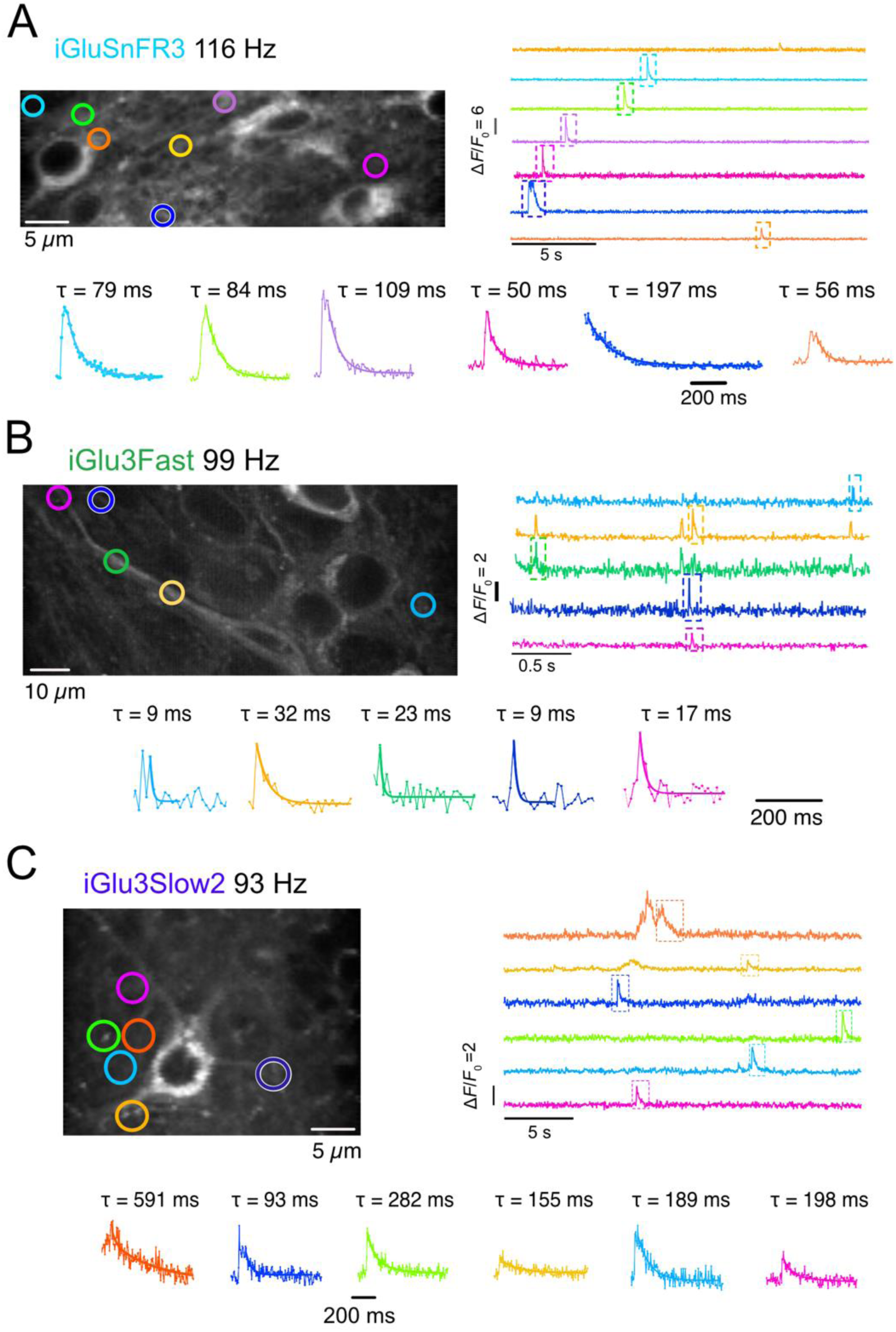
Representative recordings of spontaneous glutamate transients in mouse organotypic hippocampal slices expressing (**A**) iGluSnFR3, (**B**) iGlu3Fast or (**C**) iGlu3Slow2. 940 nm excitation, CA1 region, DIV 16. Micrographs show Z-projection of the standard deviation of the field of view of acquisition. The coloured ROIs (circles) correspond to the traces on the right, which were acquired at 115 Hz, 99 Hz, and 93 Hz, respectively and are presented unfiltered. Lower inset panels show single events (indicated on each trace by boxes with a dashed line). Traces are plotted unfiltered, as individual points, overlaid with the corresponding fitted single exponential decays.

**Figure 9.**
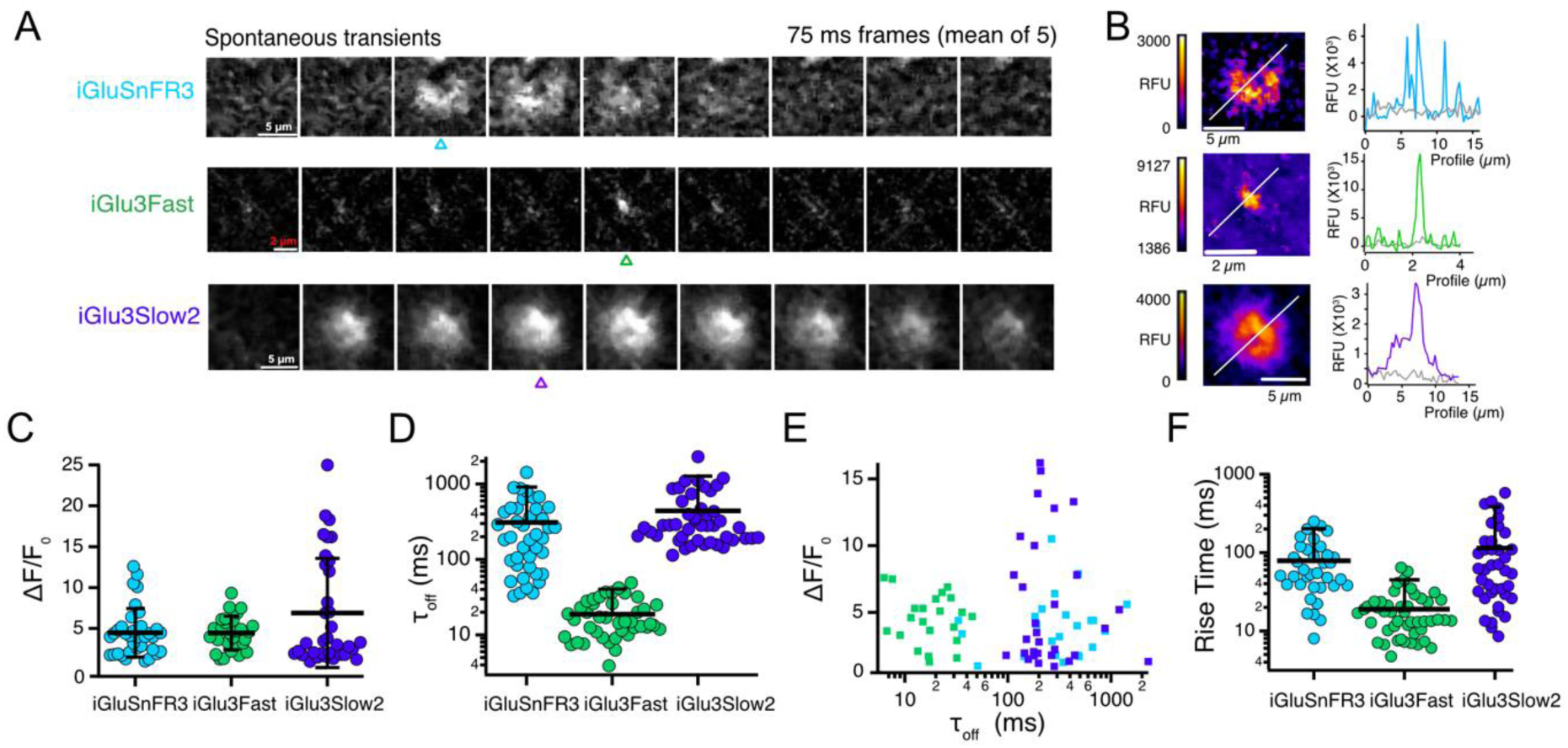
Summary of iGluSnFR3 variant spontaneous responses in organotypic slice. (**A**) Time-lapse of spontaneous release highlighting the different properties of events for iGluSnFR3, iGlu3Fast and iGlu3Slow2. Each micrograph shows the average of 5 successive frames acquired at 66-Hz original frame rate (75 ms per averaged micrograph). The iGlu3Fast event was observed in a dendrite, imaged with 63x objective. iGluSnFR3 and iGlu3Slow2 were imaged with a 20x objective. (**B)** Background-subtracted, smoothed heatmaps of the peak responses (from the frames marked with the coloured triangles) with corresponding line profiles. Colour scale bars are displayed for each heatmap. **(C)** Maximum dynamic range (Δ*F/F*_0 max_) by trace. The peak fluorescent response in each trace corresponded on average to 4.5 ± 2.9 (iGluSnFR3, *n* = 35), 4.4 ± 2 (iGlu3Fast, *n* = 28), 6.9 ± 6.7 (iGlu3Slow2, *n* = 34), respectively. **(D)** Decay time constants for spontaneous glutamate transients *τ*_off_ (per event). The data fitted well to single exponential. Mean decay τ values were 311± 303 ms (iGluSnFR3, *n* = 41), 19 ± 11 ms (iGlu3Fast, *n* = 42), 440 ± 420 ms (iGlu3Slow2, *n* = 45), respectively. Error bars represent standard deviation of the mean. (**E**) Scatter plot of the relation between the peak fluorescent dynamic range (Δ*F/F*_0 max_) and the decay time constant (*τ*_off_). For the estimation of dynamic range, several well-isolated transients were selected from each trace. **(F)** 10-90% rise times of spontaneous glutamate transients in hippocampal slice. The rise time (10-90%) extracted from the raw traces was 79 ± 62 ms (iGluSnFR3, *n* = 36), 19 ± 13 ms (iGlu3Fast *n* = 47) and 110 ± 140 ms (iGlu3slow1, *n* = 39). Error bars represent standard deviation of the mean.

iGlu3Fast showed sparse signals in organotypic slice, characterized by dimmer baseline expression and a lower apparent level of spontaneous activity, reflecting its decreased affinity for glutamate compared to iGluSnFR3. Careful examination of time series showed highly-localized fast transients (**Figure 8B**) with 10-90% rise times averaging 19 ms and mean decay times of 18 ms (**Figure 9D&F**). The decay of iGlu3Fast signals is comparable to the fast AMPA current at synapses, indeed similar to the ‘shutter speed’ of acquisition in our experiments (around 10 ms). The fastest decays that we could measure for iGlu3Fast (acquired at 193 Hz) were in the 10 ms range (**Supplementary Figure 3**, **Supplementary Movie 1**), whilst still having an excellent signal above baseline. These measurements confirm that the measured decays were not limited by imaging speed.

We observed unexpected features in the performance of iGlu3Slow2 in organotypic slices (**Figure 8C**). Slices expressing iGlu3Slow2 often exhibited uneven baseline fluorescence, with some regions showing higher intensities in the absence of detectable activity, while others displayed a weaker signal. Occasionally, slowly propagating fluorescence waves were also observed. Slices expressing iGlu3Slow2 also exhibited synchronized, temporally patterned activity (∼77%; 7 out of 9 recordings, from 4 slices obtained from different animals; **Supplementary Figure 4**, **Supplementary Movie 2**). This large-scale activity, which extended beyond the field of view (∼100 µm), occurred with markedly higher prevalence in iGlu3Slow2-expressing slices compared to those expressing iGluSnFR3 where synchronized activity was observed in only ∼33% (3/9 recordings, **Supplementary Figure 4**, **Supplementary Movie 3**). Moreover, during these events, peak fluorescence signals in iGlu3Slow2-expressing slices were substantially brighter than those observed in slices expressing iGluSnFR3. Strikingly, synchronized activity was virtually absent in slices expressing iGlu3Fast (∼14%; 1/7 recordings), except for a single instance, which coincided with tissue damage due to excessive laser power.

Strikingly, time-lapse imaging of spontaneous release highlighted the distinct kinetic properties of the three variants, with iGlu3Fast standing out for its exceptional speed (**Figure 9A**). Independent of the sample rate (up to 200 Hz), transients that were limited to just a couple of frames were invariably resolved (**Figure 8B**), suggesting that the true fluorescence events were sometimes too fast to be captured in detail (about 10 ms timescale). In contrast, events reported by iGluSnFR3 and iGlu3Slow2 tended to be larger and with some degree of internal structure (**Figure 9B&C**)

For iGlu3Slow2, a subset of events had a very large dynamic range (**Figure 9C** and **Supplementary Figure 4**). Transients were somewhat faster than in patch clamp fluorometry, which could be partly due to the increased temperature (34°C compared to room temperature). There was no correlation between the dynamic range and the decay kinetics of events for any of the reporters (**Figure 9E)** but the fast-decaying events reported by iGlu3Fast clustered separately from the slower events reported by iGluSnFR3 and iGlu3Slow2, although the tissue was held in the same conditions for each sensor. Perhaps surprisingly, the fastest kinetics (including rise time) for iGlu3Slow2 were not much different from those of iGluSnFR3 (**Figure 9D&F)**, despite the much slower kinetics of iGlu3Slow2 measured *in vitro* and expressed in HEK293T cells. This observation suggests that the events being detected were limiting the rate of the iGluSnFR3 response, rather than fast events being detected as slow purely due to IgluSnFR3 kinetics.

## Discussion

The development of genetically encoded glutamate sensors pioneered by iGluSnFR (1), allows imaging neural activity by directly monitoring neurotransmission. The iGluSnFR3 generation represented a notable improvement in the fluorescence dynamic range (in our hands, from ∼1.5 to about 6.5 in HEK293T cells (**Figure 6D&E**). The faster response due to an increased *on*-rate was, however, accompanied by decay kinetics (∼100 ms) about 100-fold slower than synaptic glutamate transients (1 ms). Previous work showed iGluSnFR3 to have a large signal-to-noise ratio but kinetics were measured with stimulation or averaging over tens of time-locked trials (6). Despite ultrafast imaging, transients with decays in the seconds range were measured in organotypic slice and in zebrafish (23) when high-activity conditions (elevated potassium, or K^+^ channel block with 4-AP) were used (6).

iGlu3Fast, the S72T variant of iGluSnFR3 has a 5-fold faster decay rate than iGluSnFR3, comparable to the ultrafast decay of iGlu*_u_*, which was derived from iGluSnFR by the same mutation (3). In addition, the S72T mutation further increased the fluorescence dynamic range *in vitro* of iGluSnFR3 from 48 to 57-fold. Notably, iGlu3Fast showed an increase in affinity for aspartate compared to iGluSnFR3, in contrast with the strong reduction in glutamate affinity. This property has been exploited to develop this variant as a genetically encoded fluorescent aspartate sensor (19).

The 3.18 mM *K*_d_ for glutamate measured for iGlu3Fast in solution was reduced to 148 - 302 μM when iGlu3Fast was expressed on the plasma membrane of HEK293T cells, showing a similar effect to that previously reported for iGluSnFR and its variants (1,3,6). However, the affinity of iGlu3Fast is still much lower than the parent sensor, iGluSnFR3, and appears favourable for preferential detection of high, transient local glutamate concentrations. This property makes it ideal for measuring glutamate in real time, in or near the synaptic cleft, in the physiological range **(Figure 8A&B)**. As the duration of the fluorescence response is reduced by the faster decay, the overall photon yield for a brief event is also reduced, which in principle should hinder detection. Despite this disadvantage, overall iGlu3Fast gave similar signal strength to iGluSnFR3 in brain slices, due to its greater molecular brightness. Overall, sensors able to report synaptic glutamate dynamics with similar kinetics to the fastest neurotransmitter receptors represent a stunning new level of performance.

The utility of iGlu3Fast with greater fluorescence dynamic range and faster kinetic responses requires imaging systems capable of taking advantage of the improved sensor properties. In hippocampal slice, the spontaneous signals from iGlu3Fast were relatively sparse, and could often only be identified *post hoc*. In contrast, the slow decay variants iGlu3Slow1 and iGlu3Slow2 offer slower kinetics and stable signals with comparatively low demands on imaging speed and sensitivity, because of the concomitant boost in photon yield for each event detected. As well as isolated, compact (<5 µm) transients, probably corresponding to vesicular release at synapses, iGlu3Slow2 reports large scale, slow events that do not appear to arise from simple summation of synaptic release events. To our knowledge, such events have not previously been widely reported. Slow glutamate “plumes” were reported in a mouse model of migraine or in hypoxic conditions, but these were tightly localised events on the micron scale (24,25). iGlu3Fast did not show global glutamate events spanning the field of view, suggesting that its low affinity for glutamate tends to filter out these events. The underlying nature of these synchronized large-area glutamate events will require careful investigation in future experiments. Several phenomena could drive these signals, including “reversed” glutamate uptake as occurs during ischaemia (26) or coordinated release of glutamate through volume regulated channels as is seen during spreading depression (27). The slow events might reflect damage to the tissue, but slices appeared otherwise healthy, and those prepared in this way were used by our group and others for decades for electrophysiology and imaging experiments, under similar conditions. If indeed these glutamate events represent a pathological state, the iGlu3Slow2 reporter appears to be a reliable and straightforward way to observe this condition.

In summary, this work demonstrates the utility of ultrafast and slow iGluSnFR3 variants for resolving rapid synaptic release events and large-scale accumulation of glutamate, respectively, by two-photon microscopy, making them excellent candidates for in vivo imaging of brain activity across scales.

*Note: Since this work was submitted, a fourth generation of iGluSnFR sensors, developed under high throughput screening, was reported with similar properties to the sensors we describe here.* https://www.biorxiv.org/content/10.1101/2025.03.20.643984v1

## Acknowledgements

This work was supported by BBSRC grant BB/S003894/1 to K.T. and by the DFG under Germany’s Excellence Strategy (EXC-2049-390688087-NeuroCure) and the large equipment initiative for innovative light microscopy (INST 276/750-1-413917635) to A.J.R.P. We thank Thorsten Trimbuch and the Viral Core Facility of the Charité Berlin for assistance with virus production and Ana Sanchez-Moreno for help with slice culture.

## Author contributions

KT conceived the project, KT and AJRP designed the project and KT wrote the first draft of the manuscript. OT and HJH generated the variants, OT carried out the in vitro measurements, LRL (Erasmus, University of Barcelona), HH and TC performed affinity measurements by live cell imaging of HEK293T cells, SB carried out patch-clamp fluorometry on HEK293T cells and two photon imaging in organotypic slice. Luka Lozo is thanked for his assistance with in vitro measurements. All authors analysed data and contributed to the writing.

---

**Supplementary Figure 1.**
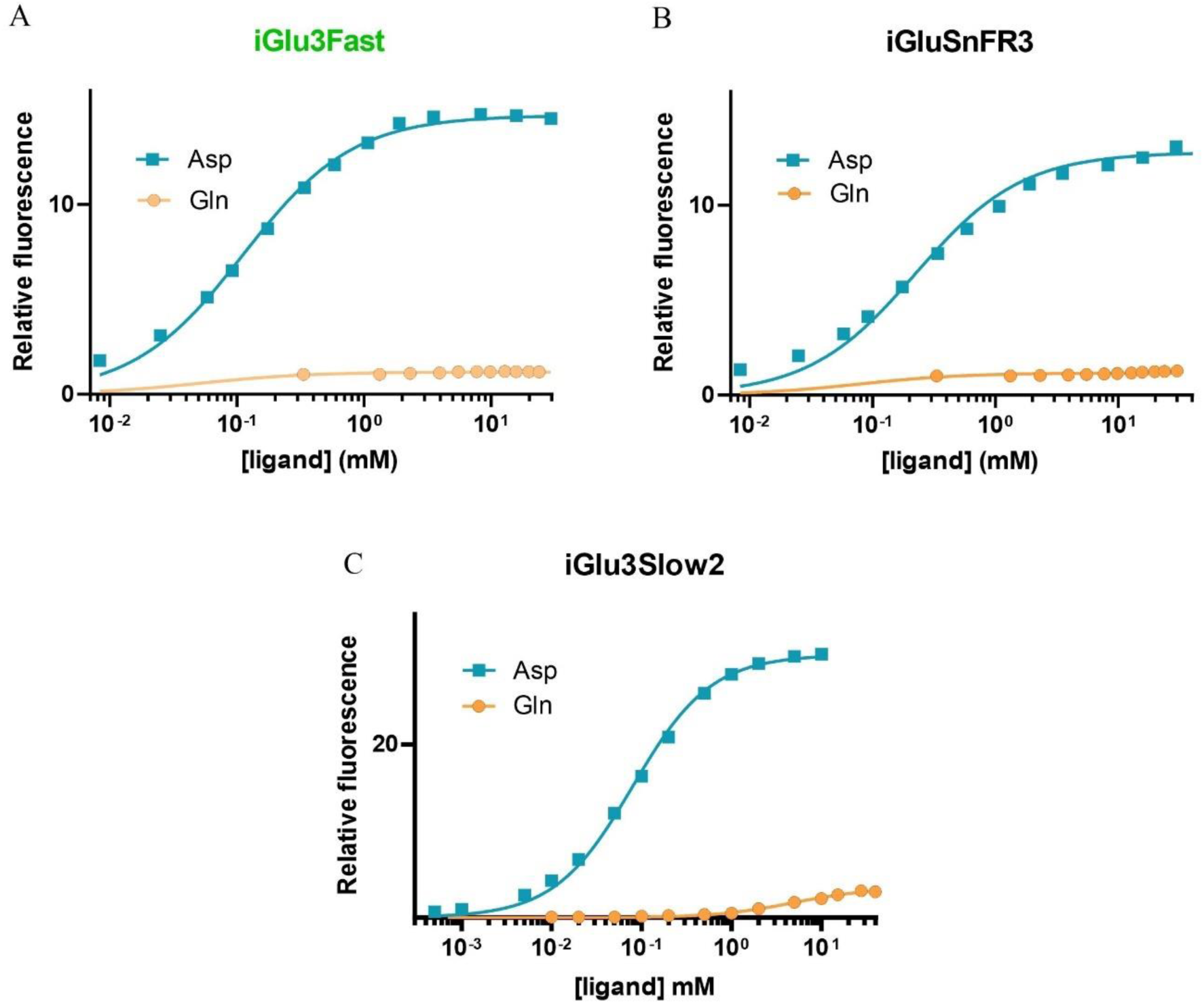
Ligand selectivity. Equilibrium titrations of (**A**) iGlu3Fast, (**B**) iGluSnFR3 and (C) iGlu3Slow2 with aspartate and glutamine, as indicated, at 20 °C.

**Supplementary Figure 2.**
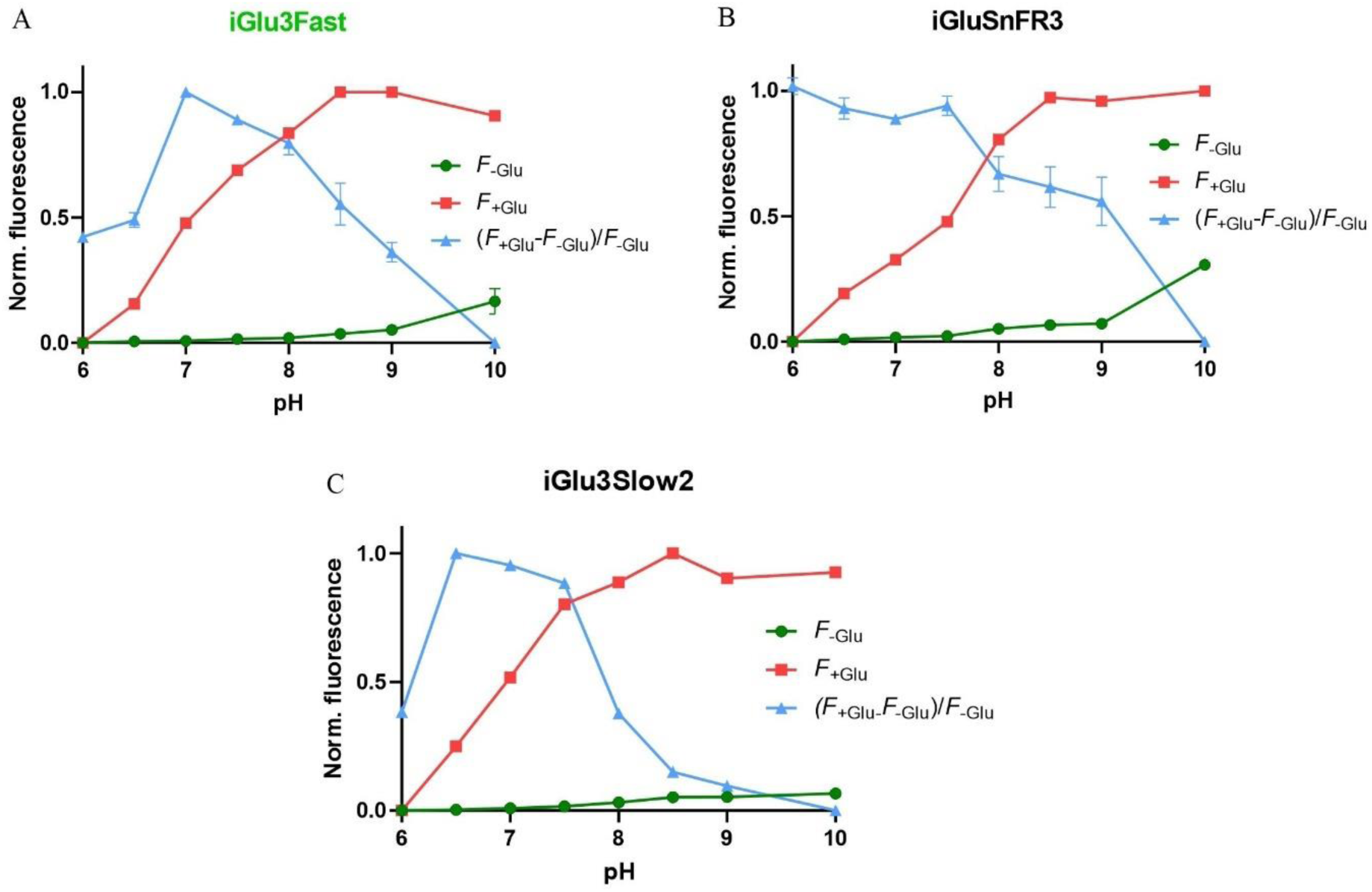
pH sensitivity and p*K*_a_ determination of (**A**) iGlu3Fast, iGluSnFR3 (**B**) and iGlu3Slow2 (**C**). Normalised fluorescence in the presence (-▪-) and absence of 30 mM glutamate (-·-).

**Supplementary Figure 3.**
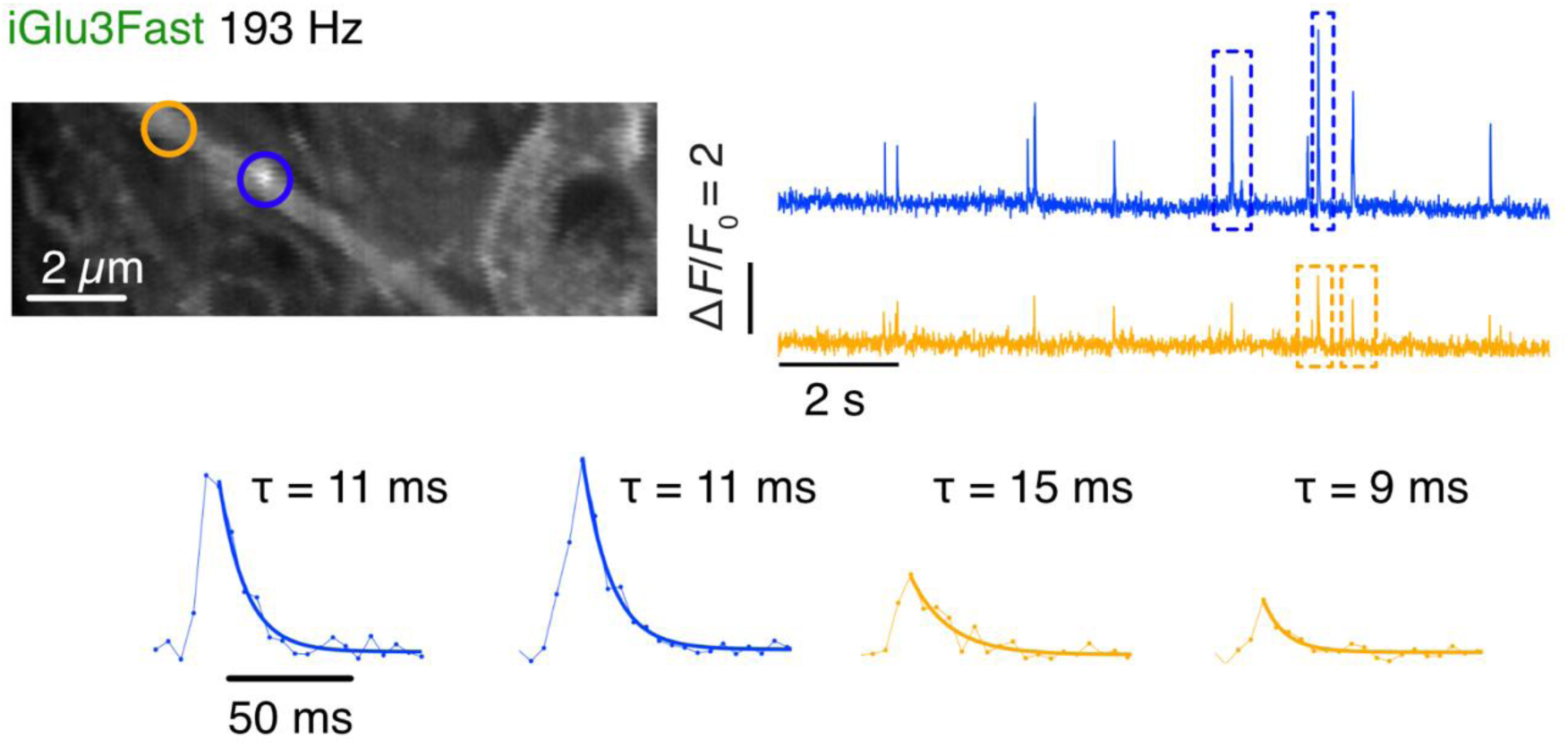
Sensitive detection of ultrafast glutamate transients in mouse organotypic slice by iGlu3Fast. In a similar region of interest as in **Figure 8B**, substantial spontaneous glutamate transients with decay constants in the 10 ms range were recorded. Micrograph is the z-projection of standard deviation of the movie (see **Supplementary Movie 1**). These transients were localised to the dendritic regions marked with circles. Lower panels show single exponential fits to single raw data traces without filtering.

**Supplementary Figure 4.**
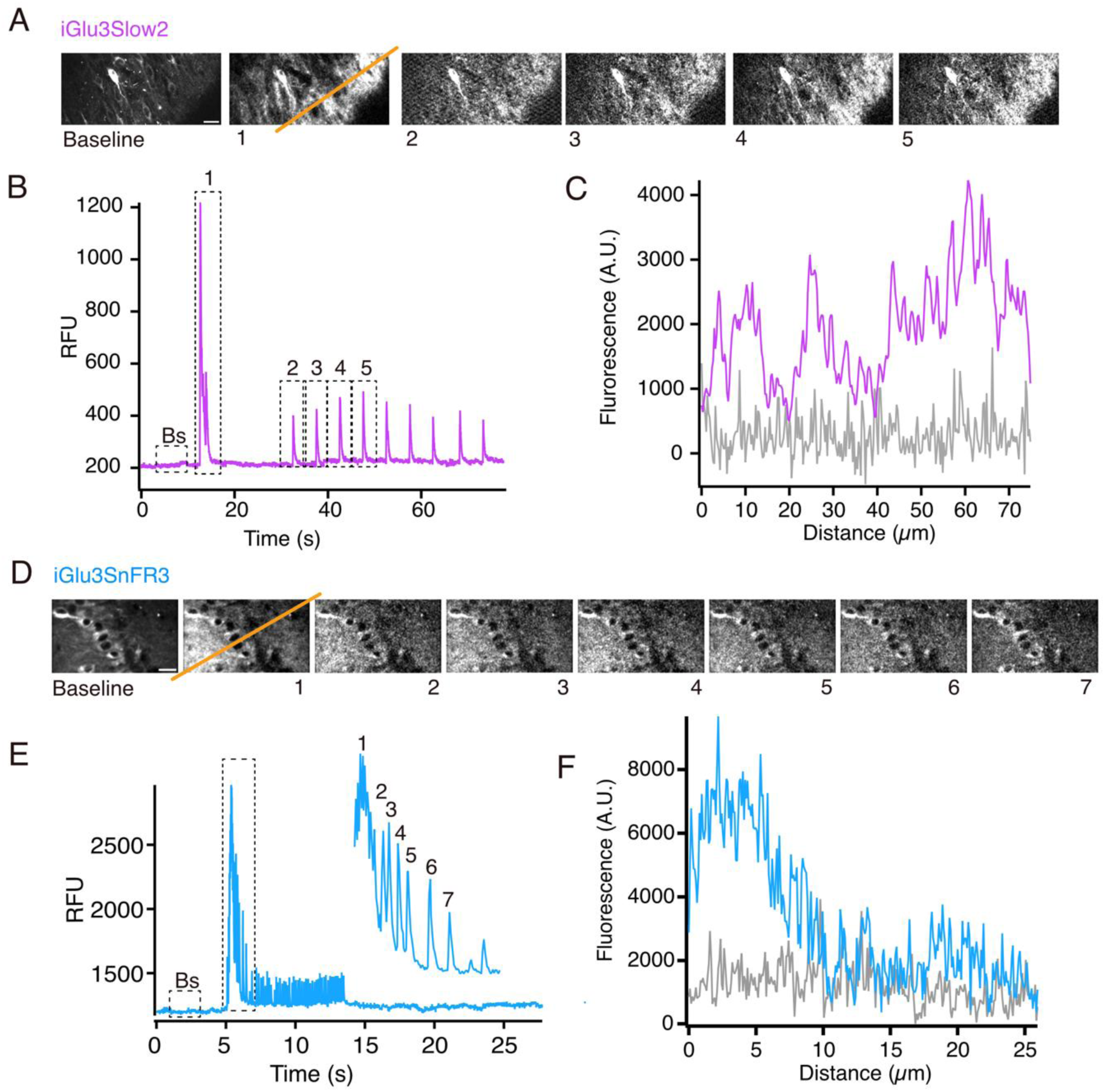
Large scale glutamate transients in mouse organotypic slice. For iGlu3Slow2 (A-C) and iGluSnFR3 (D-F), slow synchronized, temporally tuned activity was regularly seen that extended beyond the field of view. See **Supplementary Movies 2** and **3**). (A & D) Each micrograph displays the peak of fluorescent signal observed in successive numbered coordinated glutamate transients during the traces (parts B and E) which are the mean of the whole field fluorescence against time. Scale bars in each “Baseline” panel are 10 μm. (B & E) Each trace plots the full-field mean fluorescence signal. The baseline fluorescence images in A and D are taken from the part of the trace marked “Bs”. (C & F) For each line profile (diagonal orange lines superimposed on fluorescence micrographs #1, parts A and D, respectively), the fluorescence signal averaged over 5 s is displayed (iGlu3Slow2, purple and iGluSnFR3, light blue). The resting baseline fluorescence along the same profile is shown for comparison in grey.

**Supplementary Table 1.** Molecular brightness measurements for iGlu3Fast, iGluSnFR3, iGlu3Slow1 and iGlu3Slow 2.

| Variant | iGlu3Fast |  | iGluSnFR3 |  | iGlu3Slow1 |  | iGlu3Slow2 |  |
| --- | --- | --- | --- | --- | --- | --- | --- | --- |
|  | - Glu | + Glu | - Glu | + Glu | - Glu | + Glu | - Glu | + Glu |
| $\epsilon_0(492 \text{ nm})$<br>$\text{M}^{-1}\text{cm}^{-1}$ | 1873 | 27924 | 1065 | 14389 | 3337 | 38275 | 539 | 35754 |
| $\phi$ | 0.25 | 0.94 | 0.26 | 0.93 | 0.18 | 0.78 | 0.65 | 0.64 |
| Brightness<br>($\epsilon_0 \times \phi$)<br>$\text{mM}^{-1}\text{cm}^{-1}$ | 0.57 | 17.1 | 0.47 | 13.4 | 0.62 | 29.7 | 0.35 | 35.7 |
| $F_{\text{max}}/F_{\text{min}}$ | 1 | 57 | 1 | 48 | 1 | 48 | 1 | 65 |

$\phi$ , Quantum yield; $\epsilon_0$ extinction coefficient.

**Supplementary Table 2.**
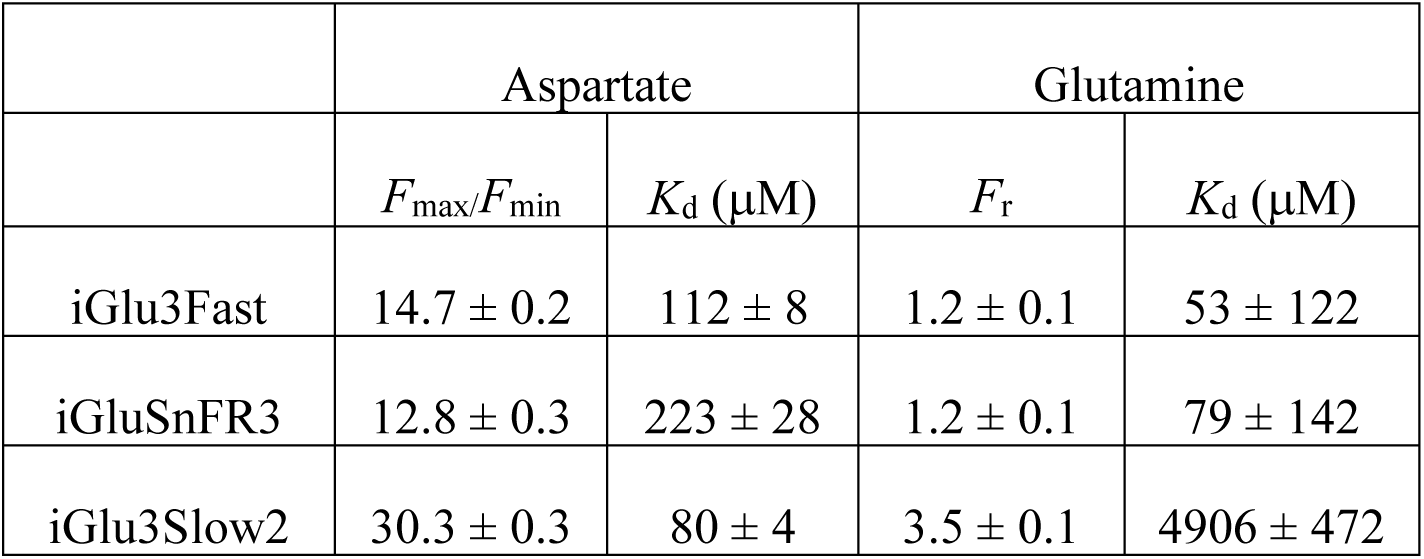
Ligand selectivity of iGlu3Fast and iGlu3Slow2 compared to iGluSnFR3.

## Legend for Supplementary Movies

**Supplementary Movie 1**

Ultrafast transients from the iGlu3-Fast reporter. Two-photon imaging at 940 nm in mouse organotypic slice. Movie was rendered following 5x temporal averaging and a 0.8-pixel radius median filter with the Green Fire Blue Look Up Table (LUT). Movie playback is at natural speed. Animated trace of the original recording at the position indicated by the white arrow, unprocessed, is shown below the movie. Source data is the same recording as in **Supplementary Figure 3**.

**Supplementary Movie 2**

Whole field spontaneous activity revealed by the iGlu3-Slow2 reporter. Two-photon imaging at 940 nm in mouse organotypic slice. Movie was rendered following 10x temporal averaging and a 0.8-pixel radius median filter with the KTZ IndiGlo LUT (https://github.com/kwolbachia/KTZ-LUTs). Playback is at 2x normal speed. Animated trace of the original recording (mean of the entire field of view), unprocessed is shown below the movie. Source data is the same recording as in **Supplementary Figure 4**.

**Supplementary Movie 3**

Whole field spontaneous activity revealed by the iGluSnFR reporter. Two-photon imaging at 940 nm in mouse organotypic slice. Movie was rendered following 10x temporal averaging and a 0.8 pixel radius median filter with the KTZ CampFire LUT. Playback is at 2x normal speed. Animated trace of the original recording (mean of the entire field of view), unprocessed, is shown below the movie. Source data is the same recording as in **Supplementary Figure 4**.

